# Menstrual cycle-driven hormone concentrations co-fluctuate with white and grey matter architecture changes across the whole brain

**DOI:** 10.1101/2023.10.09.561616

**Authors:** Elizabeth J. Rizor, Viktoriya Babenko, Neil M. Dundon, Renee Beverly-Aylwin, Alexandra Stump, Margaret Hayes, Luna Herschenfeld-Catalan, Emily G. Jacobs, Scott T. Grafton

**Author notes:** Contributed equally as first author (Elizabeth J. Rizor, Viktoriya Babenko).

## Abstract

Cyclic fluctuations in hypothalamic-pituitary-gonadal axis (HPG-axis) hormones exert powerful behavioral, structural, and functional effects through actions on the mammalian central nervous system. Yet, very little is known about how these fluctuations alter the structural nodes and information highways of the human brain. In a study of 30 naturally cycling women, we employed multidimensional diffusion and T_1_-weighted imaging during three estimated menstrual cycle phases (menses, ovulation, mid-luteal) to investigate whether HPG-axis hormone concentrations co-fluctuate with alterations in white matter (WM) microstructure, cortical thickness (CT), and brain volume. Across the whole brain, 17β-estradiol and luteinizing hormone (LH) concentrations were directly proportional to diffusion anisotropy (μFA), while follicle-stimulating hormone (FSH) was directly proportional to cortical thickness. Within several individual regions, FSH and progesterone demonstrated opposing associations with mean diffusivity and cortical thickness. These regions mainly reside within the temporal and occipital lobes, with functional implications for the limbic and visual systems. Lastly, progesterone was associated with increased tissue and decreased CSF volumes, with total brain volume remaining unchanged. These results are the first to report simultaneous brain-wide changes in human WM microstructure and cortical thickness coinciding with menstrual cycle-driven hormone rhythms. Strong brain-hormone interaction effects may not be limited to classically known HPG-axis receptor-dense regions.

## Introduction

On average, people who menstruate experience about 450 menstrual cycles throughout the lifespan (Chavez-MacGregor et al., 2008). Driving these cycles are rhythmic fluctuations in hypothalamic-pituitary-gonadal axis (HPG-axis) hormones such as sex steroids (17β-estradiol and progesterone) and pituitary gonadotropins (luteinizing hormone [LH] and follicle stimulating hormone [FSH]). The cycle begins with menses, signaling the start of the follicular phase and the gradual rise of 17β-estradiol concentrations stimulated by FSH (Baird, 1987). Just before ovulation (release of a mature egg), three “ovulatory” hormones (17β-estradiol, LH, and FSH) reach peak values; post-ovulation, the luteal phase begins, during which progesterone peaks and 17β-estradiol remains high (Reed & Carr, 2000). Fluctuations in concentrations of these hormones have been found to coincide with significant variation of typical cognition and brain function (Beltz & Moser, 2020; Chung et al., 2016; Pritschet et al., 2020). In addition, fluctuations of these hormones can induce or exacerbate neurological and psychiatric symptomatology (Dreher et al., 2007; Handy et al., 2022; Little & Zahn, 1974; Moos et al., 1969; Österlund & Hurd, 2001; Woolley & Schwartzkroin, 1998). Due to the widespread presence of gonadal hormone receptors in the mammalian brain (Barth et al., 2015; Lei et al., 1993; McEwen & Milner, 2017; Ramirez & Zheng, 1996), HPG-axis hormones exhibit powerful neuromodulatory effects that influence synaptic plasticity (Adams et al., 2001; Blair et al., 2015; Brann et al., 2007; Haraguchi et al., 2012), dendritic spine density (Woolley & McEwen, 1992, 1993), and myelination (Atkinson et al., 2019; Baulieu & Schumacher, 2000; Curry & Heim, 1966). Yet, despite the functional and cellular importance of these hormones, neuroimaging research dedicated to our understanding of the human brain as an endocrine organ makes up a very small percentage of all neuroimaging studies (Taylor et al., 2021).

In human neuroscience, the majority of published work has documented how menstrual cycle-driven hormonal fluctuations influence patterns of brain communication while participants are engaged in a cognitive/affective task (Bannbers et al., 2012; Dan et al., 2019; Jacobs & D’Esposito, 2011; Jacobs et al., 2015; Petersen et al., 2018; Pletzer et al., 2019) or at rest (“functional connectivity”; De Filippi et al., 2021; Fitzgerald et al., 2020; Greenwell et al., 2023; Hidalgo-Lopez et al., 2021; Lisofsky et al., 2015; Mueller et al., 2021; Petersen et al., 2014, 2019; Pletzer et al., 2016; Pritschet et al., 2020; Syan et al., 2017). Far less is known about hormonal influences on the anatomical highways and nodes that allow for such communication to occur (Dubol et al., 2021). Broadly speaking, these “highways” are the white matter tracts that transfer information between grey matter nodes, and both highways and nodes may vary in structural properties. Studies typically assay these neuroanatomical variables non-invasively using diffusion tensor imaging (white matter microstructure) and voxel-based morphometry (grey matter volume). White matter microstructure has been found to be altered across hormonal transition periods, including puberty, oral contraceptive use, post-menopausal estrogen therapy, and gender-affirming hormone treatment (Baroncini et al., 2010; Ha et al., 2007; Herting et al., 2012; Kranz et al., 2017; Nabulsi et al., 2020; Peper et al., 2008; Rametti et al., 2012). Only a handful of studies have investigated time-varying changes in white matter microstructural properties across a natural menstrual cycle (i.e., not affected by pharmacological interventions). These studies suggest that both WM volumetric and diffusion properties are predominantly altered during the ovulatory phase of the cycle, or correlated with 17β-estradiol concentrations (Barth et al., 2016; De Bondt, Van Hecke, et al., 2013; Meeker et al., 2020; Şafak, 2019). A larger corpus of work has probed the effect of menstrual cycle stage on regional and global grey matter volume. Grey matter morphology appears to be sensitive to hormonal transition periods, including puberty, oral contraceptive use, pregnancy, menopause, and gender-affirming treatments (Herting et al., 2014; Hoekzema et al., 2017, 2022; Lisofsky et al., 2016; Mosconi et al., 2021; Petersen et al., 2015, 2021; Pol et al., 2006; Witte et al., 2010; Zeydan et al., 2019). Only a small body of studies have investigated time-varying changes in grey matter morphology across a natural menstrual cycle. Grey matter morphology tends to change in concert with ovulation and/or 17β-estradiol (Barth et al., 2016; De Bondt et al., 2016; De Bondt, Jacquemyn, et al., 2013; Franke et al., 2015; Hagemann et al., 2011; Petersen et al., 2015), although grey matter morphological alterations are also observed more broadly between the luteal and follicular cycle phases (Lisofsky et al., 2015; Meeker et al., 2020; Ossewaarde et al., 2013; Pletzer et al., 2010) and tied to circulating progesterone concentrations (Pletzer et al., 2018; Taylor et al., 2020). A recent dense-sampling study by Zsido and colleagues (2023) used ultra-high field imaging of the medial temporal lobe, and found significant volumetric changes corresponding with 17β-estradiol and progesterone concentrations, as well as their interaction (Jacobs, 2023; Zsido et al., 2023).

The above studies are, to our knowledge, the only to map time-varying changes in structural variables across a natural menstrual cycle, highlighting the relative paucity of studies in this area, particularly in the case of white-matter (WM) microstructure. Compounding the issue for WM microstructure is the measurement typically employed; WM microstructure is commonly assessed with fractional anisotropy (FA) and mean diffusivity (MD) derived from diffusion tensor images (Beaulieu, 2002; Le Bihan et al., 2001). Despite their widespread adoption for assessing WM “integrity” in clinical populations (Clark et al., 2011; Lindenberg et al., 2010; Yuh et al., 2014), FA and MD directly scale with fiber orientation dispersion, making them sensitive to the participant-specific presence of crossing, kissing, or fanning fibers (Jones & Cercignani, 2010; Vos et al., 2012). Any single voxel within much of the WM could contain an FA value drawn from a trimodal distribution based on the number of fiber crossings (Jeurissen et al., 2013; Volz et al., 2018). Additionally, measures derived from DTI are voxel-averaged and do not account for the heterogeneity of brain tissue at the sub-voxel level (Topgaard, 2019).

However, recent developments in multidimensional diffusion imaging techniques offer a means to account for complex WM fiber configurations and sub-voxel heterogeneity when imaging human WM microstructure (Topgaard, 2017). As opposed to simple voxel-averaging, q-tensor encoding (QTE) imaging allows for the estimation of parameter distributions within each voxel, providing greater disentanglement of intersecting diffusion properties and offering a clearer picture of the underlying WM microstructure (Topgaard, 2017, 2019; Westin et al., 2016). These properties include diffusion tensor “size” (degree of free, unrestricted [isotropic] diffusion and typically aligned with increased water content), “shape” (degree of directional [anisotropic] diffusion along white matter tracts and putative measure of tissue integrity), and “orientation” (degree of fiber crossing in a voxel). QTE-derived parameters include an improved estimate of mean diffusivity (here called “D_iso_”), as well as an improved estimate of FA (called “micro fractional anisotropy”, or μFA) which is robust to fiber orientation and overcomes the associated limitations of FA (Andersen et al., 2020; Ikenouchi et al., 2020; Lasič et al., 2014). The primary aim of the present work is to therefore use recent advancements in WM imaging via QTE parameters of WM to investigate whether menstrual cycle-driven HPG-axis hormone fluctuations coincide with changes in WM microstructure.

We also report HPG-axis hormone-associated changes in grey matter cortical thickness. Previous work has mainly correlated HPG-axis hormones with changes in grey matter volume obtained through voxel-based morphometry, for which results can be heavily influenced by spatial smoothing, image co-registration imperfections, and voxel-wise correction for multiple comparisons (Whitwell, 2009). On the other hand, cortical thickness modeling techniques provide greater sensitivity in aging and imaging genetics studies (Hutton et al., 2009; Winkler et al., 2010) than volumetric analyses. Additionally, cortical thickness measures mimic functional network organization and predict clinical symptomatology (He et al., 2007; Pettigrew et al., 2016; Walhout et al., 2015). Together, these imaging techniques (QTE diffusion and cortical thickness) will provide the clearest combined account to date of how HPG-axis hormones may influence the anatomical highways and nodes of the brain.

In addition, a third aim of this study is to assess HPG-axis hormone-associated changes in brain volume. No study, to our knowledge, has investigated whether hormone-related changes in WM microstructure and cortical thickness coincide with HPG-axis hormone-related changes in estimates of total brain volume. A previous study identified peak GM volume and decreased cerebrospinal fluid (CSF) at ovulation (Hagemann et al., 2011); yet, the dynamic relationships between total brain volume, tissue volume, and CSF volume across the menstrual cycle are largely unknown and can provide insight into the potential mechanisms behind short-term WM and GM structural changes.

The current study addresses documented methodological concerns in the field of menstrual cycle neuroimaging by achieving direct hormone assay, whole brain analyses, and a robust sample size (Dubol et al., 2021). We extend previous work by assaying concentrations of four HPG-axis hormones (17β-estradiol, progesterone, LH, FSH) and recording QTE diffusion and T_1_-MPRAGE anatomical images from a group of 30 naturally cycling young women. In order to capture significant variation in hormone concentrations, we obtained data for each participant during three estimated menstrual cycle time-points: menses, ovulation, and the mid-luteal phase. We then employed a Bayesian framework to test if concentrations of each hormone would credibly predict within-individual WM and GM architecture, both at the whole brain and predefined region levels. Lastly, we examined whether these hormones additionally predicted changes in whole brain, tissue, and CSF volume.

## Results

We report data collected from 30 naturally cycling, young, healthy female participants (mean age = 21.73 years; range = 18-29) as part of a wider women’s health study at the University of California, Santa Barbara (Babenko, 2023). Data reported here includes concentrations of HPG-axis gonadal steroid hormones (17β-estradiol, progesterone) and pituitary gonadotropins (LH, FSH), parameters derived from q-tensor encoding (QTE) MRI and diffusion tensor distribution modeling (**Figures 1-2**; see Methods: *<u>Diffusion Parameters</u>)* that describe aspects of white matter diffusion tensor size, shape, and orientation, and cortical thickness measures derived from T_1_-MPRAGE anatomical images. For each participant, data were recorded during three experimental sessions estimated to coincide with three estimated phases of their individual menstrual cycles: menses, ovulation, and the mid-luteal phase (see Methods: *<u>Menstrual Cycle Tracking Procedures</u>*). To capture HPG-axis hormone variation across the menstrual cycle, study sessions were scheduled via individualized menstrual cycle tracking. For our study population, group mean participant cycle length (average across all tracked cycles for a given participant) was 31.20 days (range = 24.44 – 43.82). All participants provided written informed consent for study procedures approved by the University of California, Santa Barbara’s Institutional Review Board/Human Subjects Committee.

**FIGURE 1.**
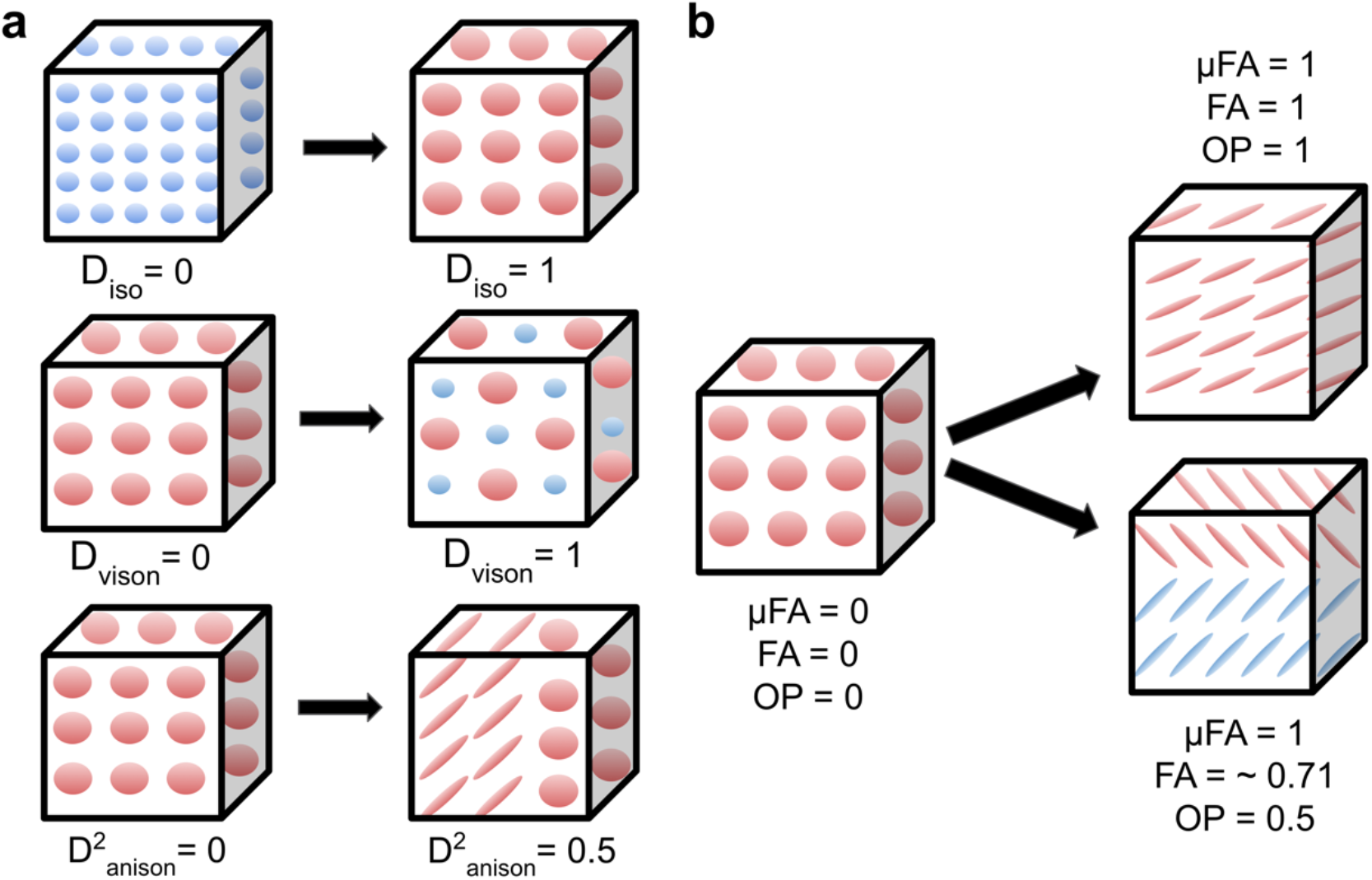
Microstructural diffusion tensor size-shape-orientation properties captured via q-tensor encoding (QTE)-imaging and diffusion tensor distribution (DTD) modeling within a voxel (cube). **1a.** Top row: An increase in Diso represents an increase in isotropic diffusion tensor size (greater isotropic diffusion is represented as larger spheres). Diso can be employed as an index of mean diffusivity. Middle row: An increase in V_ison_ represents an increase of variation in isotropic diffusion tensor size (variety of sphere size). Bottom row: An increase in D^2^_anison_ represents an increase in diffusion tensor anisotropy and a change in tensor shape (tensor shape changes from sphere to ellipse). The bottom row also depicts how DTD modeling is sensitive to microstructural variation at the sub-voxel level, such as within voxels that contain half isotropic diffusion and half anisotropic diffusion (represented here as a mixture of ellipses and spheres). **1b.** QTE-imaging and DTD modeling also allows for the estimation of other parameters describing tensor shape (conventional FA, μFA) as well as diffusion tensor orientation (OP). Left: In voxels with random orientation and isotropic diffusion (lack of parallel white matter fiber bundle), FA, μFA, and OP are all equal to 0. Right top: In voxels with a single white matter fiber bundle travelling in one direction, FA, μFA, and OP are all equal to 1. Right bottom: In voxels with crossing fibers (OP<1), FA deceases in value while μFA remains equal to 1 (robust to crossing fibers). After DTD modeling, we calculated white matter regional summary values of DTDs that describe facets of diffusion tensor size, shape, and orientation. Figure design is based on Topgaard, 2019.

**FIGURE 2.**
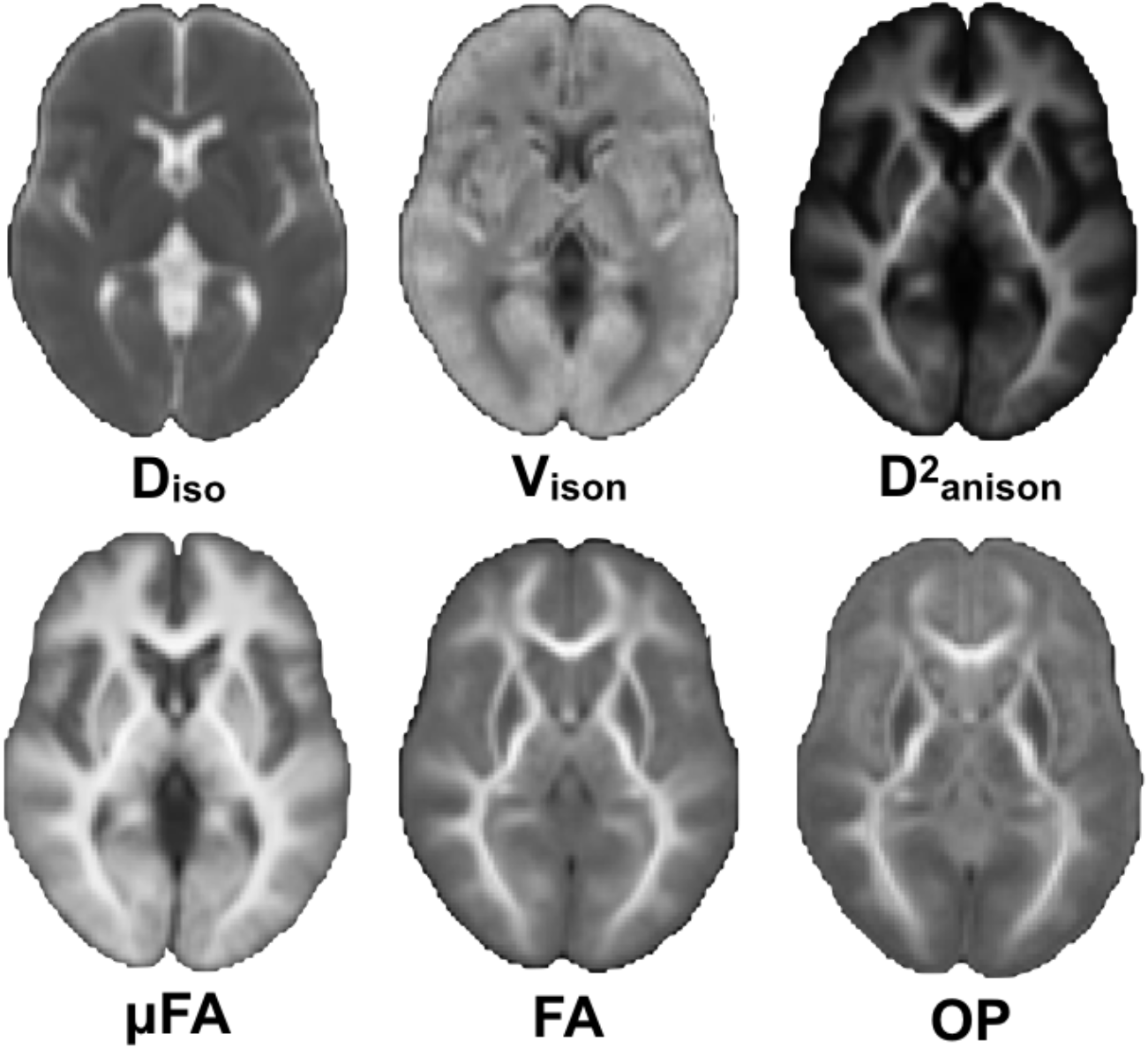
QTE-derived diffusion tensor size-shape-orientation parameter maps averaged across all 30 participants. Top row: Diso (left) is greater (brighter) in regions of greater isotropic (free water) diffusion, such as within ventricles. In contrast, V_ison_ (middle) is lower (darker) in the ventricles due to the uniform lack of restriction on isotropic diffusion, leading to lower variety in diffusion tensor size. D^2^_anison_ (right) is greater in the white matter due to increased anisotropy, similarly to measures of fractional anisotropy. Bottom row: μFA (left) remains high in white matter regions with with crossing fibers (lower OP values, right), while FA (middle) decreases in regions where OP is low.

### HPG-axis hormone concentrations

In the first step of our analyses, we verified that HPG-axis hormone concentrations varied across the three experimental sessions coinciding with estimated menstrual cycle phases (menses, ovulation, and mid-luteal). A summary of median HPG-axis hormone concentration values by estimated phase can be found in **Table 1**. One way-repeated measures ANOVAs confirmed that there were significant (p < 0.05) estimated phase on HPG-axis hormone concentration effects for 17β-estradiol (F = 40.20, p < 0.001, η^2^ = 0.47), progesterone (F = 65.93, p<0.001, η^2^ = 0.61), LH (F = 38.65, p < 0.001, η^2^ = 0.47), and FSH (F = 47.69, p < 0.001, η^2^ = 0.48).

**TABLE 1:** Median values of group serum HPG hormone concentrations by session type.

| Session Type | 17β-estradiol (pg/mL) | Progesterone (ng/mL) | LH (mIU/mL) | FSH (mIU/mL) |
| --- | --- | --- | --- | --- |
| Menses | 26.0 (17.25, 31.0) | 0.09 (0.07, 0.18) | 4.64 (3.06, 6.26) | 5.88 (4.56, 7.09) |
| Ovulation | 200.0 (125.0, 267.5) | 0.65 (0.53, 1.06) | 43.69 (21.21, 59.96) | 9.81 (6.15, 13.54) |
| Mid-luteal | 125.0 (83.25,167.5) | 11.0 (5.48, 17.25) | 4.67 (2.33, 9.72) | 2.70 (2.10, 3.94) |

Values in table are reported as median (Q1,Q3).

Post-hoc paired Wilcoxon signed-rank tests found that within-subject group-level differences in HPG-axis hormone concentration levels were consistent with known patterns of HPG-axis hormone fluctuations across a typical menstrual cycle in young, naturally cycling women (Reed & Carr, 2000; Stricker et al., 2006). Significance was defined according to a Bonferroni-adjusted p value of p < 0.0167 (p = 0.05/3 sessions). Overall, we observed a group-level pattern of moderate levels of FSH and relatively low concentrations of 17β-estradiol, progesterone, and LH during menses sessions, relatively high concentrations of 17β-estradiol, LH, and FSH during ovulation sessions, and relatively high concentrations of progesterone/moderate concentrations of 17β-estradiol during mid-luteal sessions. Specifically, 17β-estradiol concentrations (Figure 3a) were significantly greater during the ovulation and mid-luteal sessions when compared to menses (p < 0.0001), while the difference between ovulation and mid-luteal sessions did not pass the Bonferroni-adjusted p value (p = 0.028).

**FIGURE 3.**
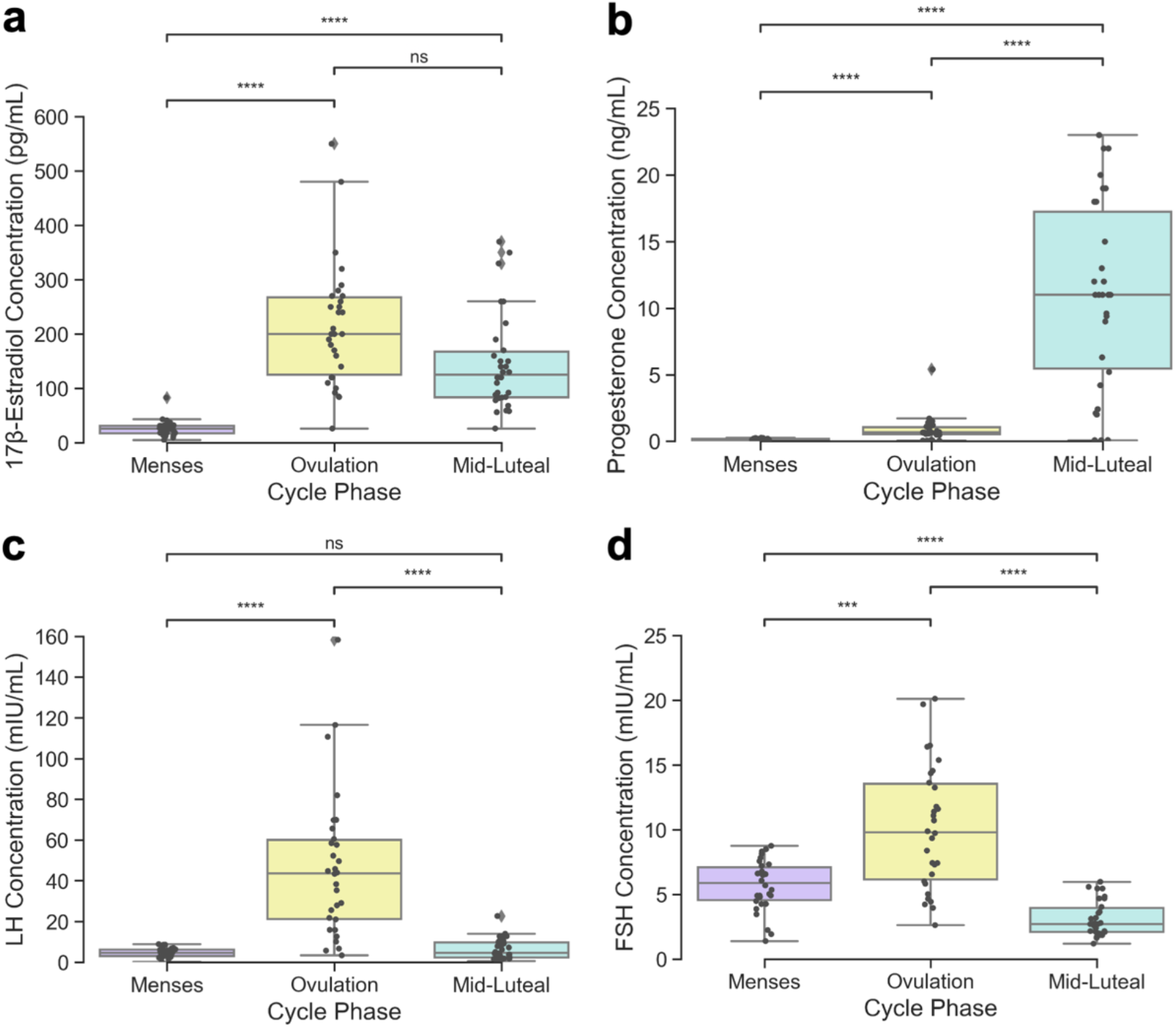
Variation in HPG-axis hormone concentrations across three experimental scanning sessions coinciding with estimated menstrual cycle phases (N=30). One way repeated-measures ANOVAs found significant estimated phase on hormone concentration effects for all four hormones studied (p < 0.001). Significant differences in within-subject hormone concentrations between sessions were determined by Wilcoxon signed-rank tests and defined with a Bonferroni-adjusted p < 0.0167. Legend: *** indicates p < 0.001, **** indicates p < 0.0001, ns indicates no significance found. Grey points indicate individual participant values.

Progesterone concentrations (Figure 3b) were significantly greater during the mid-luteal session when compared to ovulation and menses, as well as during the ovulation session when compared to menses (p < 0.0001). LH concentrations (Figure 3c) were greater during the ovulation session when compared to both menses and mid-luteal sessions (p < 0.0001); menses and mid-luteal sessions did not significantly differ (p = 0.62). Similarly, FSH concentrations (Figure 3d) were greater during the ovulation session when compared to both menses (p < 0.001) and mid-luteal (p < 0.0001) sessions, while menses was found to have greater concentrations when compared to the mid-luteal session (p < 0.0001).

### HPG-axis hormone and white matter microstructure relationships

Next, we tested if HPG-axis hormone concentrations (17β-estradiol, progesterone, LH, FSH) predict 6 white-matter (WM) diffusion parameters derived from q-tensor encoding (QTE) imaging (Figures 1-2) that describe aspects of diffusion tensor size (mean isotropic diffusivity size [D_iso_], variation in isotropic diffusivity size [V_ison_]), shape (mean squared anisotropy [D^2^_anison_], micro fractional anisotropy [μFA], fractional anisotropy [FA]), and orientation (orientation parameter [OP]). To do this, we tested the within-subject relationship between each hormone and mean ROI WM diffusion parameter values with hierarchical Bayesian regression models that could assess relationships both within 34 pre-specified WM ROIs and across the whole brain (aggregation of all ROIs) in a single model. Note that tables report unadjusted regression coefficients (β weights), while relationships were only considered credible (positive or negative) if an estimate of effect size (i.e., shrinkage-adjusted coefficients) for whole brain (D) or region-specific relationships (d_r_) exceeded defined regions of practical equivalence (ROPEs; D: [-0.06 to 0.06], d_r_: [-0.35 to 0.35]; see Methods: *<u>Statistical Analyses</u>*).

*Whole Brain:* Beginning with credible relationships across the whole brain (**Table 2**), hierarchical Bayesian regression models found that whole brain relationships between WM diffusion parameters and hormones differed depending on whether we were examining predictions of diffusion tensor size, shape, or orientation.

**TABLE 2:** Whole brain beta weights and credibility tests for all white matter diffusion hierarchical Bayesian regression models.

| Relation | $\beta_1$ | D | HDI of D |
| --- | --- | --- | --- |
| D<sub>iso</sub>-17β-estradiol | -0.055 | -0.123 | [-0.22, -0.031] |
| **D<sub>iso</sub>-Progesterone** | **0.258\*<sup>†</sup>** | **0.593** | **[0.46, 0.732]** |
| **D<sub>iso</sub>-LH** | **-0.211\*** | **-0.516** | **[-0.648, -0.397]** |
| **D<sub>iso</sub>-FSH** | **-0.299\*<sup>†</sup>** | **-0.796** | **[-1.001, -0.619]** |
| **V<sub>ison</sub>-17β-estradiol** | **0.132\*** | **0.311** | **[0.209, 0.419]** |
| V<sub>ison</sub>-Progesterone | -0.043 | -0.074 | [-0.155, 0.003] |
| **V<sub>ison</sub>-LH** | **0.085\*** | **0.193** | **[0.097, 0.288]** |
| V<sub>ison</sub>-FSH | 0.056 | 0.105 | [0.025, 0.189] |
| **D<sup>2</sup><sub>anison</sub>-17β-estradiol** | **0.182\*** | **0.381** | **[0.289, 0.48]** |
| **D<sup>2</sup><sub>anison</sub>-Progesterone** | **0.090\*** | **0.162** | **[0.081, 0.247]** |
| **D<sup>2</sup><sub>anison</sub>-LH** | **0.115\*** | **0.293** | **[0.182, 0.405]** |
| D<sup>2</sup><sub>anison</sub>-FSH | 0 | 0.001 | [-0.093, 0.095] |
| **μFA-17β-estradiol** | **0.145\*** | **0.305** | **[0.211, 0.4]** |
| μFA-Progesterone | 0 | 0 | [-0.082, 0.082] |
| **μFA-LH** | **0.111\*** | **0.261** | **[0.157, 0.364]** |
| μFA-FSH | 0.065 | 0.132 | [0.04, 0.219] |
| FA-17β-estradiol | 0.052 | 0.090 | [0.012, 0.167] |
| **FA-Progesterone** | **0.105\*** | **0.219** | **[0.121, 0.322]** |
| FA-LH | 0.027 | 0.049 | [-0.031, 0.133] |
| FA-FSH | -0.045 | -0.093 | [-0.179, -0.0] |
| OP-17β-estradiol | -0.005 | -0.010 | [-0.095, 0.071] |
| OP-Progesterone | 0.043 | 0.086 | [-0.001, 0.178] |
| OP-LH | 0.030 | 0.067 | [-0.027, 0.163] |
| OP-FSH | 0.026 | 0.056 | [-0.037, 0.152] |

\* indicates a credible relation (HDI of D lies outside of the ROPE [-0.06-0.06])
<sup>†</sup> indicates credible relations at the region level (HDI of $d_r$ lies outside of the ROPE [-0.35-0.35] for at least one region). Beta weights and credibility tests for individual region D<sub>iso</sub>-FSH and D<sub>iso</sub>-Progesterone relationships are located in Supplementary Material Tables S1-S2.

With parameters related to size (**Table 2**), we first observed that D_iso_ (a measure of isotropic diffusivity size and index of mean diffusivity) was positively predicted by progesterone concentrations (β_1_ = 0.258, D = 0.593, HDI = [0.46, 0.732]), and strongly negatively predicted by LH and FSH (LH: β_1_ = -0.211, D = -0.516, HDI = [-0.648, -0.397]; FSH: β_1_ = -0.299, D = -0.796, HDI = [-1.001, -0.619]). Meanwhile V_ison_ (variation in isotropic diffusivity size) was positively predicted by 17β-estradiol and LH (17β-estradiol: β_1_ = 0.132, D = 0.311, HDI = [0.209, 0.419]; LH: β_1_ = 0.085 D = 0.193, HDI = [0.097, 0.288]). No other relationships credibly exceeded the ROPE. Overall, these results indicate that the mean size of isotropic diffusivity (mean diffusivity) was predicted opposingly by progesterone versus LH and FSH, while variation in the size of isotropic diffusivity was positively predicted solely by 17β-estradiol and LH.

Next, with parameters related to shape (**Table 2**), we first observed that D^2^_anison_ (a measure of mean squared anisotropy) was positively predicted by 17β-estradiol, LH, and progesterone (17β-estradiol: β_1_ = 0.182, D = 0.381, HDI = [0.289, 0.48]; LH: β_1_ = 0.115, D = 0.293, HDI = [0.182, 0.405], progesterone: β_1_ = 0.09, D = 0.162, HDI = [0.081, 0.247]). Meanwhile, μFA (micro fractional anisotropy) was positively predicted by 17β-estradiol and LH (17β-estradiol: β_1_ = 0.145, D = 0.305, HDI = [0.211, 0.4]; LH: β_1_ = 0.111, D = 0.261, HDI = [0.157, 0.364]). In contrast, progesterone positively predicted conventional FA (β_1_ = 0.105, D = 0.219, HDI = [0.121, 0.322]). All other relations did not credibly exceed the ROPE. Overall, these results indicate that QTE diffusion parameters that assess diffusion tensor shape (D^2^_anison_ and μFA) were mainly predicted by 17β-estradiol and LH, similar to the pattern observed above for the size parameter V_ison_. Interestingly, progesterone predicted conventional FA, another measure of diffusion tensor shape, highlighting a discrepancy between what these anisotropy parameters may represent. Lastly, with parameters related to orientation, we observed no credible associations (**Table 2**).

*Region-specific:* We next tested relationships between HPG-axis hormone concentrations and WM diffusion parameters at the region-specific level within the same hierarchical Bayesian regression models (**Table 2**). Overall, only the size parameter D_iso_ (index of mean diffusivity) was credibly predicted by hormone concentrations at the regional level (Figure 4). More specifically, concentrations of progesterone and FSH predicted D_iso_ in multiple regions, but in opposing directions. FSH (Figure 4, top row) negatively predicted D_iso_ in 17 regions; in seven of those, progesterone (Figure 4, bottom row) also positively predicted D_iso_ (corpus callosum forceps major, corpus callosum tapetum, fornix, frontal parahippocampal cingulum, optic radiation, posterior corticostriatal tract, posterior thalamic radiation). In all credible regions, region-specific relationships trended in the same direction as the respective whole brain effects (i.e., FSH: negative, progesterone: positive). No credible region-specific relations were observed between any hormones and either shape or orientation WM diffusion parameters. These results indicate that, in several overlapping regions across the brain, progesterone and FSH concentrations had opposing influence uniquely on size parameters, i.e., they predicted mean diffusivity with opposite directionalities. Further region-specific visualization of relationships between hormone estimates and D_iso_ are provided in the Supplementary Results (**Figures and Tables S1-S2**).

**FIGURE 4.**
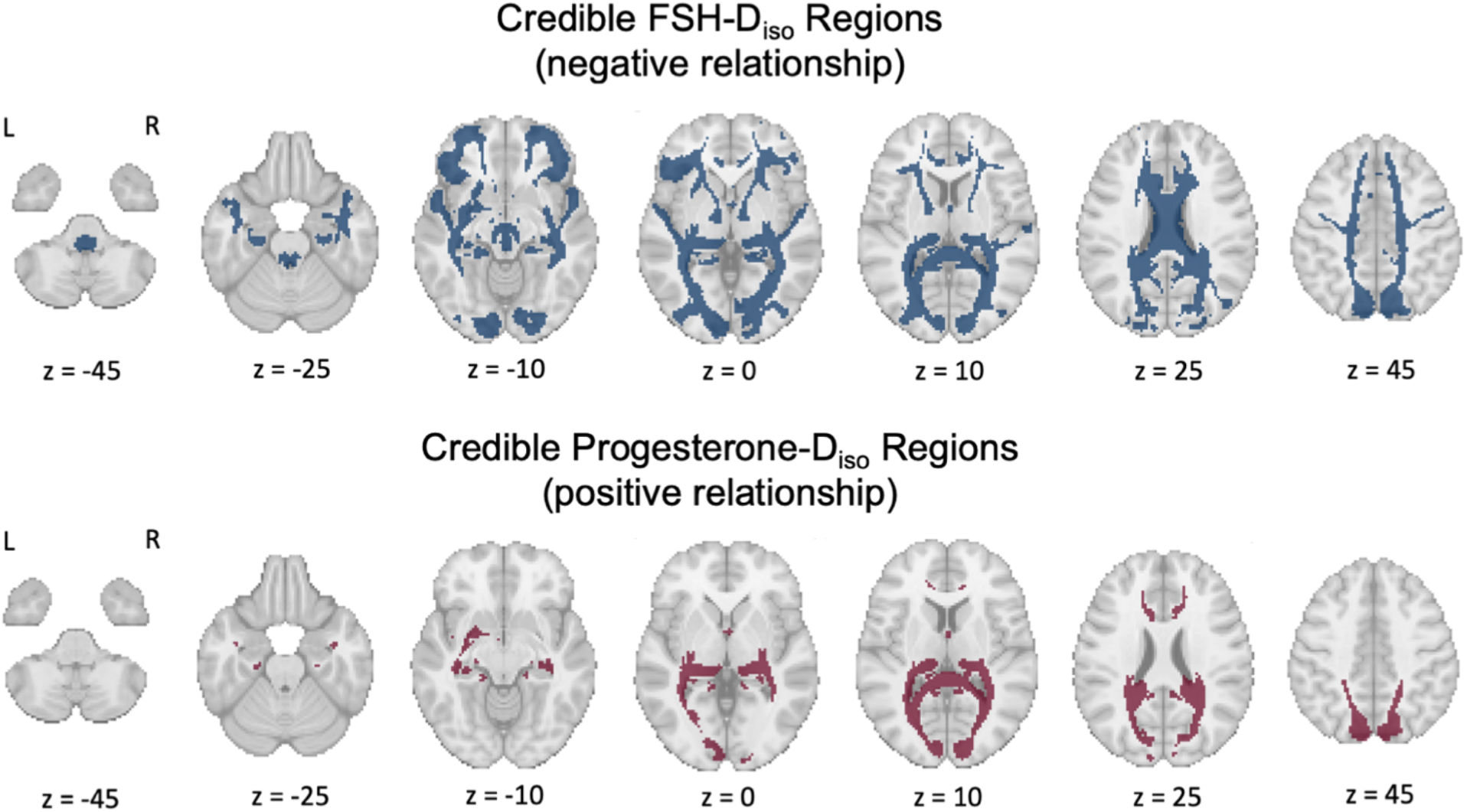
White matter regions where FSH-D_iso_ (top row) and progesterone-D_iso_ (bottom row) relationships are credible (N=30). Top row: Blue indicates regions where within-subject increases in FSH concentrations credibly predicted a decrease in mean region D_iso_ across the three sessions (negative relationship). Across the whole brain, FSH and LH concentrations credibly negatively predicted D_iso_ (LH not credible at the region-specific level). Bottom row: Red indicates regions where within-subject increases in progesterone concentrations credibly predicted an increase in mean region D_iso_ across the three sessions (positive relationship). Across the whole brain, progesterone concentrations credibly positively predicted D_iso._ Both FSH and progesterone predicted changes in D_iso_ in seven shared regions (corpus callosum forceps major, corpus callosum tapetum, fornix, frontal parahippocampal cingulum, optic radiation, posterior corticostriatal tract, posterior thalamic radiation), but in opposing directions matching their respective whole-brain trends (FSH: negative, progesterone: positive).

### HPG-axis hormone and cortical thickness relationships

We next tested whether changes in HPG-axis hormone concentrations (17β-estradiol, progesterone, LH, FSH) predict changes in mean ROI grey matter cortical thickness (CT) estimates derived from T_1_-MPRAGE images. We again employed hierarchical Bayesian regression models that could assess within-subject relationships within 31 pre-specified grey matter cortical ROIs and across the whole brain (aggregation of all ROIs). Relationships were only considered credible (positive or negative) if an estimate of effect size (i.e., shrinkage-adjusted coefficients) for whole brain (D) or region-specific relationships (d_r_), exceeded defined regions of practical equivalence (ROPEs; D: [-0.06 to 0.06], d_r_: [-0.35 to 0.35]; see Methods: *<u>Statistical Analyses</u>*).

### Whole brain

Beginning again with credible relationships across the whole brain (**Table 3**), hierarchical Bayesian regression models found that whole brain CT was uniquely predicted by FSH (β_1_ =0 .162, D = 0.395, HDI = [0.115, 0.678]). Progesterone trended toward a negative prediction of CT, but the relationship did not reach the criteria for credibility (β_1_ = -0.099, D = -0.197, HDI = [-0.453, 0.066]). We did not observe credible whole brain relationships for any of the other considered hormones.

*Region-specific:* We tested relations between hormones and CT at the region-specific level within the same hierarchical Bayesian regression models (**Table 3**). Overall, CT was predicted by both progesterone and FSH at the region-specific level.

**TABLE 3:** Whole brain beta weights and credibility tests for all cortical thickness (CT) hierarchical Bayesian regression models.

| Relation | $\beta_1$ | D | HDI of D |
| --- | --- | --- | --- |
| CT-$17\beta$-estradiol | 0.065 | 0.191 | [0.059, 0.325] |
| **CT-Progesterone** | **-0.099<sup>†</sup>** | **-0.197** | **[-0.453, 0.066]** |
| CT-LH | 0.071 | 0.198 | [0.006, 0.401] |
| **CT-FSH** | **0.162<sup>\*†</sup>** | **0.395** | **[0.115, 0.678]** |

\* indicates a credible whole brain relation (HDI of D lies outside of the ROPE [-0.06-0.06])
<sup>†</sup> indicates credible relations at the region level (HDI of $d_r$ lies outside of the ROPE [-0.35-0.35] for at least one region). Beta weights and credibility tests for individual region CT-FSH and CT-Progesterone relationships are located in Supplementary Material Tables S3-S4.

More specifically, FSH credibly positively predicted CT in eight regions (Figure 5, top row). Progesterone also credibly predicted CT in eight regions (Figure 5, bottom row), but the relationship between progesterone and CT associations did not always follow its negative whole brain trend. Progesterone negatively predicted CT in six regions, and positively predicted CT in two regions.

**FIGURE 5.**
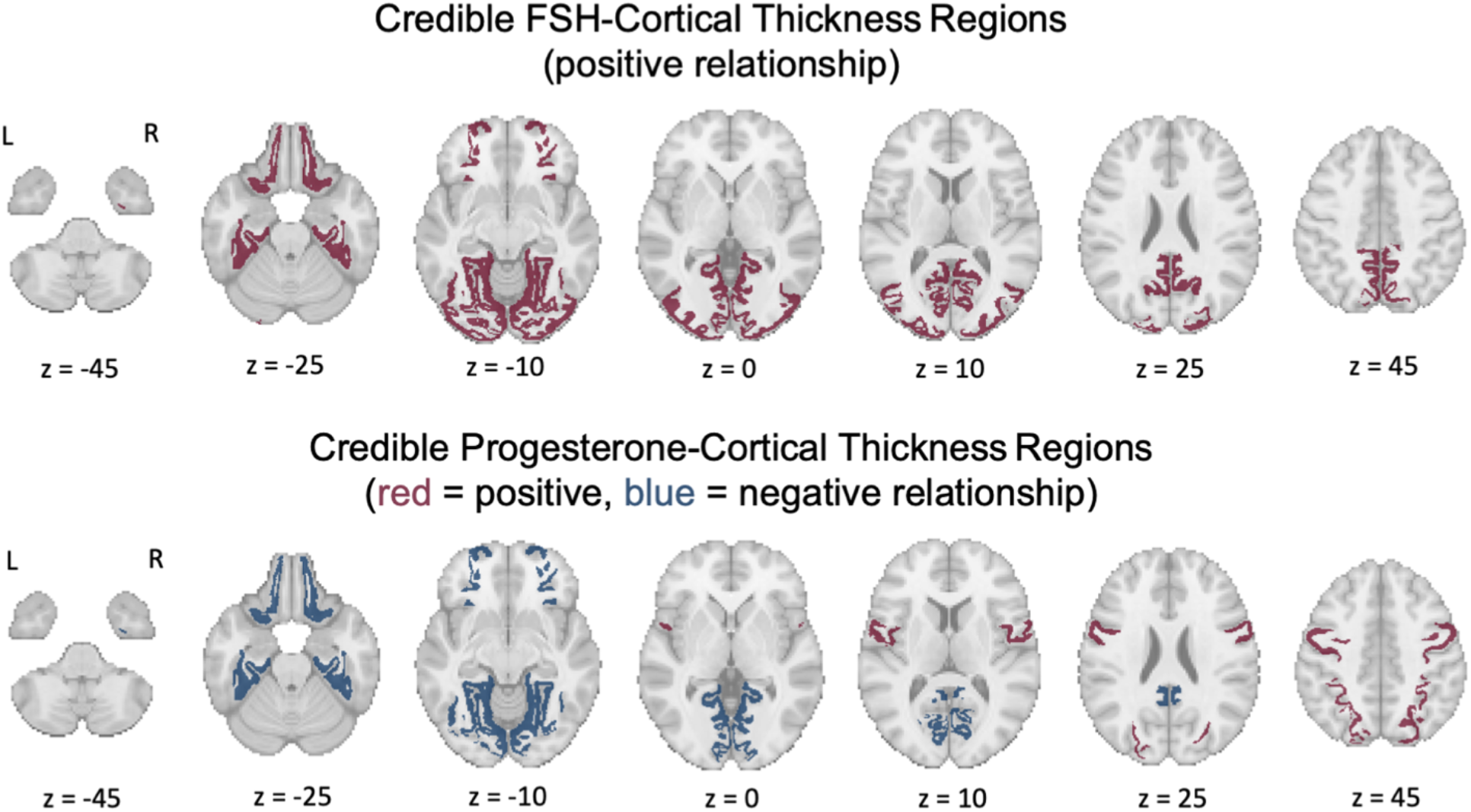
Cortical regions where FSH-CT (top row) and progesterone-CT (bottom row) relationships are credible (N=30). Top row: Red indicates regions where within-subject increases in FSH concentrations credibly predicted an increase in mean region CT across the three sessions (positive relationship). These regions include the fusiform gyrus, isthmus cingulate, lateral occipital gyrus, lateral orbitofrontal gyrus, lingual gyrus, parahippocampal gyrus, pericalcarine cortex, and the precuneus. Across the whole brain, FSH concentrations credibly positively predicted CT. Bottom row: Blue indicates regions where within-subject increases in progesterone concentrations credibly predicted a decrease in mean region CT across the three sessions (negative relationship). Positive FSH-CT relationships were also credible for these areas. These regions include the fusiform gyrus, isthmus cingulate, lateral orbitofrontal gyrus, lingual gyrus, parahippocampal gyrus, and the pericalcarine cortex. Red indicates regions where progesterone credibly predicted an increase in CT (positive relationship). These regions include the precentral gyrus and the superior parietal lobule.

Both FSH and progesterone predicted CT in six shared regions (fusiform gyrus, isthmus cingulate, lateral orbitofrontal gyrus, lingual gyrus, parahippocampal gyrus, and pericalcarine cortex); in each of these regions with shared FSH and progesterone influences over CT, relationships trended in opposite directions from each other. We did not observe any credible region-specific relationships between CT and either 17β-estradiol or LH. Based on these results, we observed that FSH and progesterone were associated with CT in opposite directions within several overlapping regions, similarly to the region-specific pattern observed for the white matter parameter, D_iso_ (mean diffusivity). However, unlike what was found in the white matter, region-specific relationships between progesterone and CT sometimes deviated from the relationships seen at the whole brain level. Further region-specific visualization of relationships between hormone estimates and CT are provided in the Supplementary Results (**Figures and Tables S3-S4**).

### HPG-axis hormone and brain volume relationships

We lastly tested whether changes in HPG-axis hormone concentrations (17β-estradiol, progesterone, LH, FSH) predict within-subject changes in total brain volume, tissue volume, and cerebrospinal fluid volume (**Table 4**). To do this, we again employed hierarchical Bayesian regression models. Note that in this case, there was no layer to the hierarchical model associated with individual brain regions, such that each variable here relates to a whole brain measure of a volumetric area. These whole brain relationships were considered credible (positive or negative) if an estimate of effect size (i.e., shrinkage-adjusted coefficients, or D) exceeded a defined region of practical equivalence which is equivalent to that of a single region in the above models (-0.35 to 0.35).

**TABLE 4:** Beta weights and credibility tests for all brain volume hierarchical Bayesian regression models.

| Relation | $\beta_1$ | D | HDI of D |
| --- | --- | --- | --- |
| Total Brain Volume-17β-estradiol | 0.111 | 0.341 | [-0.774, 2.332] |
| Total Brain Volume-Progesterone | -0.053 | -0.101 | [-0.625, 0.434] |
| Total Brain Volume-LH | 0.081 | 0.173 | [-0.471, 0.86] |
| Total Brain Volume-FSH | 0.066 | 0.167 | [-0.655, 1.239] |
| Tissue Volume-17β-estradiol | 0.279 | 2.203 | [-0.461, 20.52] |
| **Tissue Volume-Progesterone** | **0.660\*** | **2.616** | **[0.607, 15.845]** |
| Tissue Volume-LH | -0.193 | -1.058 | [-10.497, 1.008] |
| Tissue Volume-FSH | -0.452 | -1.366 | [-6.281, -0.14] |
| Cerebrospinal Fluid-17β-estradiol | -0.292 | -2.631 | [-24.278, 0.535] |
| **Cerebrospinal Fluid-Progesterone** | **-0.749\*** | **-2.709** | **[-11.604, -0.903]** |
| Cerebrospinal Fluid-LH | 0.183 | 1.262 | [-1.205, 11.979] |
| Cerebrospinal Fluid-FSH | 0.484 | 1.907 | [0.18, 12.87] |

\*Indicates credible relations at the whole brain volumetric area level (HDI of D lies outside of the ROPE [-0.35-0.35] for at least one region)

Hierarchical Bayesian regression models found that progesterone credibly positively predicted tissue volume (β_1_ = 0.66, D = 2.616, HDI = [0.607, 15.845]) and negatively predicted cerebrospinal fluid volume (β_1_ = -0.749, D = -2.709, HDI = [-11.604, -0.903]); no other hormones credibly predicted tissue or cerebrospinal fluid volumes. We did not observe credible total brain volume relationships for any of the considered hormones.

## Discussion

In the current study, we tested whether menstrual cycle-driven HPG-axis hormone concentrations in 30 naturally cycling women predicted within-subject WM microstructural, GM cortical thickness (CT), and brain volume changes at the whole brain and region-specific levels. Group-level HPG-axis hormone concentrations measured during three estimated phases of the menstrual cycle (menses, ovulation, mid-luteal) were confirmed to fluctuate in accordance with expected menstrual cycle values (Stricker et al., 2006). 17β-estradiol, LH, and FSH (hormones that peak during ovulation) were associated with increased whole brain WM microstructural anisotropy (μFA), whole brain variation in isotropic diffusion tensor size, and whole brain and region-specific CT (FSH only), as well as decreased whole brain and region-specific isotropic diffusion (mean diffusivity). In contrast, progesterone (which peaks during the luteal phase) was associated with increased mean diffusivity at both the whole brain and region-specific levels, increased and decreased region-specific CT, and increased tissue/decreased CSF volume.

### HPG-axis Hormones and white matter microstructure

While recent work has identified menstrual cycle-related structural changes in singular regions such as the hypothalamus, hippocampus, and fornix, the current study is the first to report widespread WM microstructural changes associated with cycle-driven hormone fluctuations (Baroncini et al., 2010; Barth et al., 2016; De Bondt, Van Hecke, et al., 2013). Our results suggest a global decrease in freely-diffusing WM water during the ovulatory window, leading to a momentary rise in anisotropic diffusion and variation in tensor size. Post-ovulation, we find a global increase in freely-diffusing WM water and tissue volume coinciding with the luteal rise of progesterone concentrations. While FSH and progesterone-associated changes in mean diffusivity were credible at the region-specific level, HPG-axis hormone-associated fluctuations in anisotropy and mean diffusivity identified here were overall not restricted to singular white matter bundles.

Previous work has identified 17β-estradiol-associated increases in hippocampal FA and 17β-estradiol/LH-associated decreases in fornix mean diffusivity in naturally cycling women (Barth et al., 2016; De Bondt, Van Hecke, et al., 2013). Similarly, our results suggest an ovulatory hormone (17β-estradiol and LH)-associated pattern of increased anisotropy, as well as decreased limbic system mean diffusivity. On the other hand, progesterone was found to be positively associated with mean diffusivity. Previous work has observed increased apparent diffusion coefficients (diffusivity) across the whole brain in the luteal vs. follicular phases, though the results were not statistically significant (Şafak, 2019). Yet, paradoxically, progesterone was also found to increase with FA (a ratio of anisotropy/isotropy). FA’s known sensitivity to factors outside of cell microstructure, such as fiber orientation, may explain this discrepancy (Jones & Cercignani, 2010; Volz et al., 2018). Crossing fibers are widespread in the WM across the whole brain (known as “the crossing fiber problem” in diffusion imaging; Schilling et al., 2017). While FA may provide adequate representation of WM microstructure in isolated tracts with parallel bundles, track-specific changes in FA in this study and others may be compromised when investigating across the whole brain due to potential interactions between WM structural changes and crossing fiber anatomy. Our whole brain volume observations provide context to this finding. The observed positive relationship between progesterone and tissue volume contained within a static total brain volume would cause this tissue to expand into the ventricles, leading to the observed decrease in CSF volume. This change may cause displacement of crossing fiber locations within the white matter, thus inducing spurious alterations in FA values.

### Potential mechanisms of HPG-axis hormone-associated WM microstructural changes

While the mechanism driving cycle-dependent changes in diffusion parameters across the whole brain are unknown, here we consider several putative mechanisms. Both estradiol and progesterone are implicated in the upregulation of cell myelination and myelin repair (Arevalo et al., 2010; Baulieu & Schumacher, 2000; Schumacher et al., 2012). Notably, progesterone was found to increase oligodendrocyte progenitor cell recruitment in mice, inducing axonal remyelination on the order of days (Hussain et al., 2011). Yet, the current study found that 17β-estradiol was associated with increased anisotropy (often claimed to represent increased white matter “integrity”/myelination), while progesterone was associated with increased diffusivity.

Due to the limited resolution of diffusion imaging, these parameters are sensitive to changes in several factors, including myelin content, axon density (Friedrich et al., 2020), and interstitial/extracellular fluid volume, making it difficult to pinpoint exactly which factor is contributing to these diffusion property changes.

Estradiol has also been linked with Aquaporin-4 (AQP4), a water channel ubiquitous in astrocyte end feet located on brain capillaries (Wardlaw et al., 2020). AQP4 is implicated in facilitating water movement between perivascular spaces (Virchow-Robin spaces) and the interstitial fluid, rendering its role integral for maintaining brain water and ion homeostasis, a process especially apparent during post-injury edema formation (Sun et al., 2007; Wardlaw et al., 2020). Rodent studies have identified inhibitory effects of estradiol on AQP4; for example, estradiol treatment was found to prevent AQP4 expression after rodent models of blood brain barrier (BBB) disruption and ischemic brain edema formation were induced (Shin et al., 2011; Tomás-Camardiel et al., 2005). In the latter study, ovariectomized female mice saw reversed effects (increased AQP4 induction), which were subsequently restored with estrogen receptor therapy (Tomás-Camardiel et al., 2005). In contrast, progesterone has been implicated in decreasing the inhibitory effect of estradiol on brain water accumulation (Soltani et al., 2016), a mechanism that may underlie our observed positive relation between progesterone and mean diffusivity. In addition to water movement, AQP4 may modulate gonadal steroid hormone and neurotransmitter functioning. AQP4 knockout in female mice has been found to induce subfertility, decrease estradiol and progesterone concentrations, and prevent decreases in striatal dopamine post-ovariectomy (Sun et al., 2007, 2009). This rodent evidence suggests that the ubiquitous nature of AQP4 across the brain may explain our identified global changes in brain water dynamics and have significant functional implications. Yet, while gonadal hormones have been linked to aquaporin functioning in human reproductive organs, the link between AQP4 and steroid hormones in the human brain is largely unknown and requires further study (de Oliveira et al., 2020; R.-H. He et al., 2006).

### HPG-axis hormones and cortical thickness

With regard to cortical thickness, exogenous (e.g., via oral contraceptives) and endogenous changes in HPG-axis hormone concentrations have been associated with alterations in region-specific cortical and subcortical GM morphology (Braden et al., 2017; Hoekzema et al., 2017; Lisofsky et al., 2016; Petersen et al., 2015; Taylor et al., 2020; Zsido et al., 2023). The current study is the first to report widespread cortical thickness changes directly correlated with HPG-axis hormone fluctuations across the whole brain. FSH was associated with increased CT and progesterone was associated with decreased CT within several shared regions, including the parahippocampal and fusiform gyri. Previous work has also identified increased cortical GM volume in the right fusiform and parahippocampal gyri, as well as increased left hemisphere CT, in the early follicular (i.e., preovulatory) vs. luteal phases (Meeker et al., 2020; Pletzer et al., 2010). Additionally, 17β-estradiol (typically low during the follicular phase and high during the luteal phase) has been negatively correlated with anterior cingulate cortex GM volume (De Bondt, Jacquemyn, et al., 2013). However, our finding that progesterone is associated with both increased and decreased CT suggests that the directionality of HPG-axis hormone-GM morphology relationships may vary widely across regions or even subregions. Previous studies have found increased amygdala GM volume during the luteal phase, contradicting the above-mentioned work, as well as variation in whether progesterone is positively or negatively correlated with medial temporal lobe GM volume across subregions. (Taylor et al., 2020). Based on our results, as well as the heterogeneity of HPG-axis hormone receptor locations across the brain, with known dense concentrations in the medial temporal lobe, thalamus, hypothalamus, cerebellum, brainstem, and prefrontal cortex, brain-HPG-axis hormone relationships should not be assumed to be uniform across all regions (Barth et al., 2015; Hedges et al., 2012; Österlund & Hurd, 2001).

### Potential mechanisms of HPG-axis hormone-associated cortical thickness changes

The observed cycle-associated changes in cortical thickness may be due to a variety of factors. Neuromodulatory factors beyond synaptic transmission, such as 17β-estradiol, have been shown to increase astrocytic Ca2+ and lead to neural activity-independent slow modulation of cerebral blood flow (Iadecola & Nedergaard, 2007). Additionally, arterial spin labeling MRI has identified increased frontal pole cerebral blood flow during the follicular phase, albeit with a very small sample of women (Otomo et al., 2020). Thus, the observed changes in cortical thickness may be due to hormone-driven alterations in cerebral blood flow. However, evidence suggests that cortical thickness declines in aging adults are independent from cerebral blood flow, casting doubt on this potential explanation (Chen et al., 2011). Additionally, HPG-axis hormone-associated volumetric changes in the medial temporal lobe have been found to be independent of cerebral blood flow (Zsido et al., 2023). Rodent studies have identified rapid (timescale of hours to days) estradiol and progesterone-mediated changes in hippocampal cell spine/synapse density (Woolley & McEwen, 1992, 1993), synaptic sprouting (Hara et al., 2015; Scharfman & MacLusky, 2014), and cell proliferation (Barha & Galea, 2010). Yet, these demonstrations of hormone-driven changes in neural plasticity are limited only to the hippocampus. While we identified CT changes in MTL structures such as the isthmus of the cingulate and the parahippocampal gyrus, other frontal, occipital, and parietal regions were subject to these changes as well. Due to the cytoarchitectonic diversity of the cortex, it is unclear whether these same short-term modulations would occur outside of hippocampal structures in the human brain, and whether cortical thickness even directly indicates these processes. Further study is required to determine whether these hormone-driven cellular modifications are occurring across the human cortex, how cortical thickness measures may reflect these modulations, and how they vary across cortical and subcortical regions.

### Behavioral and functional implications

Although we do not currently report functional consequences or correlates of structural brain changes, our findings may have implications for hormone-driven alterations in behavior and cognition. Notably, our results point to FSH vs. progesterone-associated opposing effects (mean diffusivity and cortical thickness) in several shared WM and GM regions. These identified regions are mainly limbic (fornix, parahippocampal gyrus, cingulate) and temporo-occipital (fusiform and lingual gyri, pericalcarine cortex, optic radiation, posterior thalamic and corticostriatal tracts). Within our sample, FSH was the only hormone that exhibited moderate values during the menses session. Thus, our observed structural changes may be due to a pre-ovulatory (moderate FSH) vs. post-ovulatory (high progesterone) “seesaw” effect, or may be directly linked to FSH and progesterone values. A wide body of evidence supports the idea that progesterone is a powerful modulator of limbic structures, rapidly altering hippocampal subfield structure (Taylor et al., 2020), enhancing rodent memory consolidation (Frick & Kim, 2018), and exhibiting paradoxical anxiolytic (Del Rio et al., 1998; Frye et al., 2006) and negative affect-promoting properties through action on GABA-A receptors (Andréen et al., 2006; Bäckström et al., 2011). Additionally, recent menstrual cycle functional imaging studies have revealed that progesterone is linked to decreased cortical and cerebellar functional network coherence, increased posterior default mode network activity, and increased functional connectivity between fronto-parietal regions and the hippocampus (Arélin et al., 2015; Fitzgerald et al., 2020; Hidalgo-Lopez et al., 2021; Pritschet et al., 2020). Along with these findings, our results support the notion that progesterone not only acts on the limbic system, but also on structural and functional networks across the whole brain. While progesterone has been more widely studied, very little is known about how endogenous FSH affects the human brain in young women. Network analyses of a single woman’s menstrual cycle, the only menstrual cycle study to date linking FSH and brain function, found that pituitary gonadotropin (LH and FSH) concentrations were coupled with network community frequency (Greenwell et al., 2023). In human and animal aging research, both FSH and FSH receptor gene polymorphisms have been associated with altered risk for developing Alzheimer’s Disease and cognitive decline (Corbo et al., 2011; Mao et al., 2022; Xiong et al., 2022). Higher FSH to 17β-estradiol ratio levels in postmenopausal women have been identified as a biomarker of cognitive impairment (Hestiantoro et al., 2017). Thus, circulating levels of FSH are associated with long-term cognitive outcomes, but it is unclear whether FSH significantly affects behavior and cognition on a short timescale, such as across the menstrual cycle. While we did not directly link changes in WM microstructure or CT to behavioral outcomes in the present study, menstrual cycle-driven changes in amygdala GM volume have been directly correlated with stress-induced negative affect (Ossewaarde et al., 2013). Another study found that women with dysmenorrhea (painful uterine cramping during menstruation) experienced enhanced menses-associated hypertropic and atropic GM volume changes in pain-related regions when compared to controls (Tu et al., 2013). Due to this, it can be inferred that our reported widespread alterations in WM and GM structure may have significant behavioral, cognitive, and clinical consequences across functional brain networks.

Finally, our results have implications for understanding how HPG-axis hormones provide neuroprotective effects in the face of conditions such as traumatic brain injury and stroke. Ample rodent evidence supports the neuroprotective effects of both estradiol (Arevalo et al., 2010) and progesterone (Brotfain et al., 2016), which have been shown to protect against both brain edema (Naderi et al., 2015; O’Connor et al., 2005; Soltani et al., 2016) and ischemic damage (Gerstner et al., 2009; L. He et al., 2014), as well as induce post-injury cell proliferation (Barha et al., 2011). Despite this, human sex difference studies often report worse brain injury outcomes for women (Gupte et al., 2019). Due to the apparent lack of HPG-axis hormone diffusion imaging studies, our understanding of how these neuroprotective mechanisms may occur in humans is very poor. As an initial step, our results demonstrate that HPG-axis hormones may be significant modulators of whole brain water dynamics and WM cell architecture. Future work is needed to directly link HPG-axis hormone concentrations with brain injury outcomes and identify the implications of these WM structural changes for women’s health.

### Limitations and future considerations

While the current study has significant strengths due to its within-subject design and reliable brain measures, several limitations exist. Data were only sampled at three time points/participant, providing sparse sampling to our regression model and leaving out associations that occur during the other days of the menstrual cycle. We estimated scheduling of the ovulation and luteal sessions through at-home ovulation tests, which can be positive across multiple days, or, for some individuals, never reach the “positive” threshold as defined by the test manufacturer. Because of this, ovulation and mid-luteal sessions may not have captured the exact ovulation or mid-luteal day for all participants. Despite this, significant fluctuations in HPG-axis hormones were captured, allowing us to achieve the primary aim of associating brain measures with HPG-axis hormone concentrations (as opposed to cycle phase). Future dense-sampling studies are needed to fill these information gaps. The correlational nature of this work means that true causality between HPG-axis hormones and brain structure could not be established. Alterations in brain structural measures may be due to non-hormone factors that coincide with menstrual cycle rhythms. While the current study aimed to address this by assessing brain-HPG-axis hormone (as opposed to cycle phase) relationships, future population studies with pharmacological hormone suppression are needed to establish causality. Additionally, while the current work has a larger sample size than many previous menstrual cycle studies, larger consortium studies are warranted to truly establish brain-hormone associations at the population level. All participants were adults under the age of 30 at time of study, but menstrual cycle-driven brain-hormone associations may evolve throughout the lifespan. Further study in younger and older age groups is needed to better understand this relationship. The cortical thickness changes described here are based on standard T_1_-MPRAGE images, which may miss more localized/subregional changes that we are not able to detect. Further study of brain-hormone relationships at the subregion level is warranted. Finally, an analysis consisting of 40 separate Bayesian models may pose a risk for susceptibility to Type 1 error. However, our imposition of a ROPE based on null-generative models serves as a conservative protection for multiple comparisons.

## Conclusions

HPG-axis hormones (17β-estradiol, progesterone, LH, FSH) were associated with significant modulations of white matter microstructure and cortical thickness. Naturally cycling young women experience widespread brain structural changes in concert with HPG-axis hormone fluctuations. While modulation of the medial temporal lobe is more established, structural and functional networks across the brain should be considered in HPG-axis hormone research. Investigation of brain-hormone relationships across networks is necessary to understand human nervous system functioning on a daily basis, during hormone transition periods, and across the human lifespan.

## Methods

### Participants

A total of 30 naturally cycling, young, healthy female participants (mean age = 21.73 years; range = 18-29) completed all study recruitment, pre-screening, and protocol procedures. Study inclusion criteria required that participants be between the ages of 18-30, have not used any hormonal or implant birth control within three months prior to onset of study involvement, self-reported no plans to begin birth control or become pregnant within the upcoming year, and have a self-reported relatively regular (21-40 day length) menstrual cycle. Exclusionary criteria included contraindications to MRI (non-removable metal, incompatible medical devices, hearing loss/tinnitus, claustrophobia), medical history of cardiovascular, neurological, or psychiatric conditions, or current use of prescription medications such as antidepressants or anxiolytics (over-the-counter pain relievers and allergy medications were accepted). All participants provided written informed consent for study procedures approved by the University of California, Santa Barbara’s Institutional Review Board/Human Subjects Committee.

Participants were recruited via digital flyers sent to the University of California, Santa Barbara community. To assess eligibility, participants completed the following: a pre-screening video call to ensure understanding of study procedures, the UCSB Brain Imaging Center’s MRI screening form, and a comprehensive health questionnaire, which assessed participant demographics, history of substance usage, and mental and reproductive health topics, for which they received $10 compensation if deemed eligible. Of a total sample of 46 eligible participants, 30 completed all 3 experimental sessions involving MRI scans and blood draws. Of these 30 participants, 22 completed at least one month of researcher-supervised menses and ovulation test tracking prior to the menstrual cycle containing their initial session. Due to study timing constraints, the remaining 8 participants began cycle-tracking within the same cycle as their initial session. Participants were paid $60 per MRI session, and those who completed a full three cycles of pre-initial session remote cycle tracking earned an additional $50, totaling $190-240/participant.

### Menstrual Cycle Tracking Procedures

Researchers engaged in cycle tracking in order to, as closely as possible, schedule experimental sessions that would match the three phases of participants’ individual cycles. To begin, participants were asked to self-report to researchers their menses start and end dates, as well as share any previous cycle tracking data they may have collected for personal use. Researchers then calculated average cycle lengths from these reports, which were used to predict future scheduling of sessions. Cycle length information was updated as new menses tracking data was collected. Menses sessions were scheduled based on a combination of self-reported menses onset, self-reported menses symptom onset, and ovulation test results.

In order to schedule ovulation and luteal sessions, participants were given 40 mL disposable plastic urine cups and ovulation testing strips, which measure urine levels of LH (Easy@Home, Premom). Experimenters informed participants of their predicted “fertile-window” (on average within a range of 3-15 days depending on prediction certainty), during which ovulation was likely to occur. Participants were asked to complete ovulation self-tests and send a clear photo of results by 8pm during all “fertile-window” days. Using the Premom mobile application, experimenters obtained a ratio of LH-to-control line darkness, for which any result > 0.80 was considered to be positive for ovulation. If a participant’s ovulation test results never produced an LH-to-control line exceeding 0.80, their ovulation test was considered to be positive according to their individual peak value determined from their history of study cycle tracking. After completing cycle tracking, experimenters informed participants of their potential ovulation session dates, during which they were asked to send in their ovulation test results by 9am. If their ovulation test result ratios were > 0.80 (or at the level of their previous individually-determined peak), they were asked to come in that day to complete study procedures. If the participant was unable to be scheduled that same day for any reason, their tracking data was logged and the session was postponed to their following cycle.

Mid-luteal sessions were scheduled based on both average cycle length and ovulation testing data. Using individualized cycle prediction data, experimenters scheduled mid-luteal sessions during the predicted mid-point between ovulation and start of next menses. If a participant could not be scheduled within their predicted menses, ovulation, or mid-luteal windows for that cycle, then experimenters postponed the session to the next cycle. If a participant received a SARS-CoV-2 vaccine/booster or diagnosis, or reported ingestion of emergency contraceptives, they waited at least one complete menstrual cycle (menses-to-menses) prior to returning for any additional sessions.

### Experimental Protocol

Participants underwent three experimental MRI sessions each, which were scheduled to coincide with three estimated phases of their menstrual cycle: menses (based on self-report), ovulation (based on ovulation test results), and mid-luteal phase (estimated to be mid-way between ovulation window and predicted start of menses). Order of sessions was counterbalanced such that 15 participants began with the menses session, 14 participants with the ovulation session, and 1 began with the mid-luteal session. Sessions lasted for 3 hours, typically took place during the hours of 11am-4pm, and did not all necessarily occur within the same monthly cycle. Participants were instructed to maintain their typical daily routines on session days. Upon arrival for their session, experimenters screened the participant for SARS-CoV-2 according to the University of California, Santa Barbara’s Office of Research COVID safety guidelines. Then, participants were instructed to change into scrubs, underwent a blood draw completed by a licensed phlebotomist (see *Blood Sample Acquisition and Processing*), and completed a 1-hour MRI scanning protocol (see *Magnetic Resonance Imaging Acquisition and Processing*).

### Blood Sample Acquisition and Processing

In order to assess serum levels of the gonadal hormones 17β-estradiol and progesterone, as well as the pituitary gonadotropins luteinizing hormone (LH) and follicle stimulating hormone (FSH), a licensed phlebotomist collected a blood sample (< 8.5 – 10 cc) from each participant during each session (3 samples/participant). The phlebotomist used a BD Diagnostics vacutainer push button to start an intravenous line (hand or forearm), and then used a 10mL vacutainer SST tube (BD Diagnostics) to collect the sample. After collection, the sample was allowed 30 minutes to clot at room temperature, and was then centrifuged (2100 RPM for 10 minutes). From the centrifuged samples, experimenters aliquoted 1mL of serum into three 2mL microtubes, which were subsequently stored in a −80° freezer until 2 of the microtubes (per participant) were shipped for processing at the Endocrine Technologies Core (ETC) at the Oregon National Primate Research Center (ONPRC, Beaverton, OR). The third microtube was stored as a backup in case any damage occurred to the samples during shipment for testing.

Gonadal steroid hormone concentrations were obtained using ultra-high performance liquid chromatography-heated electrospray ionization-tandem triple quadrupole mass spectrometry (LCMS/MS) on a Shimadzu Nexera-LCMS-8060 instrument (Kyoto, Japan). The dynamic range for both 17β-estradiol and progesterone was 0.002 to 20 ng/ml, with the following lower quantification limits, intra-assay variations, and accuracies, respectively: 0.002 ng/ml, 2.1%, and 100.9% for 17β-estradiol, and 0.010 ng/ml, 12.3%, and 106.3% for progesterone. Pituitary gonadotropin concentrations were obtained using a Roche cobas e411 automated clinical immunoassay platform (Roche Diagnostics, Indianapolis, IN). The assay range for both LH and FSH was 0.1 – 200 mIU/ml, while the intra-and inter-assay coefficients of variation (CVs) were respectively 2.3% and 2.4% for n = 2 LH assays, and 0.9% and 1.0% for n = 2 FSH assays.

### Magnetic Resonance Imaging Acquisition and Processing

Following each session’s blood draw, participants underwent magnetic-resonance imaging (MRI) in a Siemens 3T Prisma scanner with a 64-channel phased-array head/neck coil. First, high-resolution T_1_-weighted magnetization prepared rapid gradient echo (MPRAGE) anatomical scans were acquired (TR = 2500 ms, TE = 2.22 ms, FOV = 241 mm, T_1_ = 851 ms, flip angle = 7°, with 0.9 mm^3^ voxel size). Following the anatomical scan, a series of 4 spherical b-tensor (b = 0, 100 – 500, 1000, 1500 s/mm^2^; 3 diffusion directions) and 4 linear b-tensor (b = 500, 1000, 1500, 2000 s/mm^2^; 6, 10, 16, 30 diffusion directions) q-space trajectory encoding (QTE) diffusion sequences (Martin et al., 2020) were collected (TR = 6308 ms, TE = 80 ms, diffusion gradient amplitude = 80.0 mT/m, FOV = 230 mm, flip angle = 90°, 2.0 x 2.0 mm^2^ in-plane resolution, 4.0 mm slice thickness, iPAT factor = 2).

MRI preprocessing was conducted with Advanced Normalization Tools (ANTs; Avants et al., 2011) and MATLAB (The MathWorks Inc, 2018). First, T_1_-weighted anatomical data were skull-stripped with antsBrainExtraction.sh. In order to create individualized participant white matter (WM) and grey matter (GM) tissue masks, we first segmented the skull-stripped anatomical data using the ANTsPy (https://github.com/ANTsX/ANTsPy) kmeans_segmentation function, which outputted probability maps of participant WM, GM, and cerebrospinal fluid (CSF). We then binarized these probability maps (all values > 0 = 1) to obtain WM, GM, and CSF tissue masks.

Second, for each participant, we calculated ROI-specific mean values of 6 QTE-derived WM diffusion parameters, which describe facets of diffusion tensor size, shape, and orientation. To do this, we first used an open source, MATLAB pipeline for multidimensional diffusion MRI (https://github.com/markus-nilsson/md-dmri) to complete motion and eddy current correction of QTE data, as well as estimation of voxel-wise brain maps of size-shape-orientation diffusion tensor distributions (DTDs) using the “dtd” method (Nilsson et al., 2018; Topgaard, 2019). Then, using ANTs, we spatially normalized participant DTD data and HCP1065 Population-Averaged

Tractography Atlas probabilistic WM region masks (thresholded at ≥ 50%; Yeh, 2022) to their participant-specific anatomical space, where all further calculations were conducted. Next, we utilized the participant-specific WM tissue masks to extract white matter voxels from DTD data, which was then segmented into 64 regions of interest (ROIs) using the HCP1065 atlas masks. Then, for each ROI, we calculated mean values of parameters describing diffusion tensor size (average isotropic diffusivity [D_iso_] and normalized variation in isotropic diffusivity [V_ison_]), tensor shape (normalized mean squared anisotropy [D^2^_anison_], micro-fractional anisotropy [μFA], and fractional anisotropy [FA]), and tensor orientation (fiber orientation parameter [OP]; for descriptions of each, see *<u>Diffusion Parameters</u>*). Finally, because we had no *a priori* hypotheses for hemisphere-specific effects, we averaged homologous ROI mean values across hemispheres, leading to a total of 34 bilateral WM region values per diffusion parameter per participant session.

Third, for each participant, we calculated cortical and subcortical ROI-specific mean values of cortical thickness (CT). We used anatomical as opposed to diffusion imaging to obtain cortical thickness due to the known limitations of diffusion imaging for detecting neurite density (Lampinen et al., 2019). To do this, we first utilized participant-specific GM tissue masks to extract grey matter voxels from their respective anatomical images. Then, we used the ANTsPy DiReCT algorithm to estimate CT across the grey matter. DiReCT has been shown to have good scan-rescan repeatability and outperform FreeSurfer in prediction of CT measures (Tustison et al., 2014). In order to segment the grey matter into ROIs, we first obtained 20 Open Access Series of Imaging Studies (OASIS) young, healthy adult brains and their respective Desikan-Killiany-Tourville (DKT; 31 per hemisphere) cortical labels as defined by the Mindboggle project (Klein & Tourville, 2012). We then spatially normalized these labels to participant anatomical space, where we used antsJointLabelFusion.sh to obtain customized DKT-31 cortical region labels for each participant (Wang et al., 2013). Next, we utilized these customized participant-specific labels to segment the CT data and subsequently calculated mean CT values for each ROI in participant-specific anatomical space. Due again to a lack of *a priori* hypotheses for hemisphere-specific effects, we averaged the mean CT values of homologous right and left hemisphere ROIs. This led to a total of 31 bilateral GM region CT values per participant session.

Finally, for each participant session, we calculated total brain volume, tissue volume, and cerebrospinal fluid volume. To do this, we used ANTsPy label_geometry_measures with the previously generated kmeans segmentation masks to calculate volume (mm^3^) for the whole brain (WM+GM+CSF), brain tissue (WM+GM), and CSF (CSF mask only). This led to a total of three volume values per participant session.

### Diffusion Parameters

For each participant, we estimated 6 voxel-wise diffusion parameters describing diffusion tensor size (mean isotropic diffusivity; D_iso_, normalized variation in isotropic diffusivity; V_ison_), shape (normalized mean squared anisotropy; D^2^_anison_, fractional anisotropy; FA, micro fractional anisotropy, μFA), and orientation (orientation parameter; OP)

In order to calculate the diffusion tensor size (D_iso_ and V_ison_) and shape (D^2^_anison_) parameters as defined by Topgaard, 2019, we first estimated voxel-wise distributions of parallel (D_||_) and perpendicular (D_⊥_) component diffusivities (in 10^-8^ m^2^/s). Then, with these values, mean isotropic diffusivity (D_iso_) and normalized diffusion anisotropy (D_anison_, represented in Topgaard, 2019 as D_Δ_) were calculated (Topgaard, 2019, equation 2). We then took the expected values (E[x]) of D_iso_ and D_anison_ to obtain E[D_iso_] (from now on referred to as D_iso_), which is equivalent to mean diffusivity (MD), and E[D_anison_], which contains similar information as other measures of tensor shape/anisotropy, such as μFA (Topgaard, 201910/9/23 8:39:00 PM, equations 9 and 10). To obtain V_ison_, we calculated the variance of isotropic diffusivity (V[D_iso_]; Topgaard, 201910/9/23 8:39:00 PM, equation 11), which was then normalized by D^2^_iso_. Visualizations of D_iso_, D^2^_anison_, and V_ison_ are shown in Figure 1a.

In order to calculate the remaining parameters describing tensor shape (μFA, FA) and orientation (OP), we drew on other sources of QTE theory (Lasič et al., 2014; Westin et al., 2016). More specifically, μFA was calculated according to equation 14 in Lasič et al., 2014. The orientation dispersion parameter (OP), also known as “microscopic orientation coherence”, or the ratio of micro-to macroscopic anisotropy in a voxel, was calculated according to equation 33 in Westin et al., 2016. The orientation parameter ranges from 0 (diffusion in all directions equally, whether it be isotropic or anisotropic) to 1 (parallel diffusion in one direction). As discussed in Lasič et al., 2014 (equation 22), FA can be derived from μFA and OP; thus, participant μFA and OP data were used to calculate their respective FA maps. Both μFA and FA range from 0 (completely isotropic diffusion) to 1 (completely anisotropic diffusion); however, in voxels with multidirectional diffusion (high OP), μFA is designed to be robust to these orientation changes (Lasič et al., 2014, Figure 1b). Group-averaged diffusion parameter brain maps are shown in Figure 2.

### Statistical Analyses

All analyses were conducted using Python (version 3.7.10). We first aimed to verify that our session timing successfully captured group-level natural variation in gonadal steroid hormones (17β-estradiol, progesterone) and pituitary gonadotropins (LH, FSH) consistent with known typical menstrual cycle-related hormone fluctuations for young, naturally cycling women (Stricker et al., 2006). To do this, one way repeated-measures ANOVAs using the pingouin (0.5.3) package were conducted that examined the effect of session type (menses, ovulation, mid-luteal) on HPG-axis hormone concentration. Then, to determine specific session-level differences in HPG-axis concentrations, we conducted post-hoc paired-sample Wilcoxon signed rank tests (statannot 0.2.3), which controlled for the non-normality of underlying HPG-axis concentration distributions. Differences were considered to be significant if they met a Bonferroni-adjusted p value of p < 0.0167 (p = 0.05/3 sessions).

Second, we tested whether: 1) HPG-axis hormone concentration levels predict changes in white matter (WM) diffusion tensor size-shape-orientation parameters at the whole brain and region-specific levels, 2) HPG-axis hormone concentration levels predict changes in cortical thickness (CT) at the whole brain and region-specific levels, and 3) HPG-axis hormone concentration levels predict changes in brain volume. To test these relationships, we conducted Bayesian hierarchical regression models to assess whether participants’ HPG-axis hormone concentration values predicted their respective WM diffusion and CT measures across all three sessions. We sampled posterior distributions using No U-Turn sampling (NUTS) Hamiltonian Monte Carlo, implemented with the PyMC3 package (Salvatier et al., 2016). After tuning the sampler’s step size to an acceptance level of 0.95, posteriors were sampled in four parallel chains of 10000 samples (40000 total) with an additional initial 5000 samples per chain (tuning samples were then discarded). We required that no chain contained any divergences and that no posterior’s R value (the ratio of variance within chains to the variance of pooled chains), was greater than 1 (Vehtari et al., 2021). We then calculated highest density intervals (HDIs; the Bayesian equivalent of a confidence interval) using the default settings (i.e., 94% density) in the arviz package (Kumar et al., 2019).

In order to test the first two questions, 28 total models were run: 24 WM diffusion-hormone models (6 diffusion measures [D_iso_, V_ison_, D_anison_, μFA, FA, OP] × 4 hormones [17β-estradiol, progesterone, LH, FSH]), and 4 CT-hormone models (CT × [17β-estradiol, progesterone, LH, FSH]). We first separately log-transformed and z-score normalized and the brain measures across sessions within each region for each participant, and log-transformed and normalized the HPG-axis hormone concentrations across sessions within each participant. We then fitted a hierarchical model, with the lowest level sampling β0_n,r_ and β1_n,r_ parameter posteriors at each brain region r for each subject n. This level allowed for the testing of region-specific brain-hormone relations. The next level of the hierarchy constrained the distributions of β0_n,r_ and β1_n,r_ to be drawn from group-level Gaussian distributions, i.e., β0_n,r_∼N(μ_β0(r)_,σ_β0(r)_) and β1_n,r_∼N(μ_β1(r)_,σ_β1(r)_). Here, the group-level μ_β1(r)_ parameter posterior reflects the expected value of the brain-hormone relation at the group level within a region r, while σ_β1(r)_ reflects variation in this relation across individuals at that region, i.e., “shrinkage”. The next layer of the hierarchy accounted for whole brain effects at the group level by constraining μ_(r)_ and σ_(r)_ parameters to be drawn from hierarchical Gaussian distributions, i.e., μ_(r)_∼N(M_μ_,Σ_μ_) and σ_(r)_∼N(M_σ_,Σ_σ_). Here the group-level M_u(β1)_ parameter reflects the expected value of the brain-hormone relation across all regions at the group-level, while M_σ(β1)_ reflects the average shrinkage (i.e., variation in relation across subjects) across regions. M and Σ parameters were respectively constrained with uninformative Gaussian (μ∼N(0,1)) and half-Gaussian (σ∼halfN(1)) priors. This modeling framework allowed us to assess credible relations for both specific regions and for the whole brain within the same model.

To assess credible relations at the region level, we first computed a deterministic posterior (d_r_), which scaled the expected value by the shrinkage, i.e., μ_β1_/σ_β1_. For whole brain effects, we computed deterministic posterior (D), which was the average brain-hormone relation at the group level across regions, scaled by the average shrinkage (participants) in the relation across regions, i.e., M_u(β1)_/M_σ(β1)_. We then considered relations credible if the HDI of posteriors d_r_ or D exceeded region of practical equivalences (ROPEs; Makowski et al., 2019) determined with null-generative simulations using identical data structure (i.e., 34 for WM diffusion or 31 for CT regions, 30 participants, 3 sessions, variables drawn from a z-distribution; see Supplementary Methods). For the 24 WM diffusion-hormone models and 4 CT-hormone models that were run, these ROPEs were respectively (-0.35 to 0.35) for d_r_ posteriors and (-0.06 to 0.06) for D posteriors.

Finally, for the third question, we tested whether HPG-axis hormones would predict changes in total brain volume, tissue volume, and cerebrospinal fluid volume. In order to examine volume-hormone relationships, 12 models (3 volume measures [total brain, tissue, cerebrospinal fluid] × 4 hormones [17β-estradiol, progesterone, LH, FSH]) were run. As before, we z-score normalized the brain measures across sessions for each participant and normalized the HPG-axis hormone measures across sessions within each participant. For these volume-hormone models, “regions” are not defined as atlas ROIs, but as our three whole brain volumetric areas of interest (total brain, tissue, cerebrospinal fluid). These volume-hormone models featured nearly the same structure as the previously described WM diffusion-hormone and CT-hormone models; yet, there was no bottom layer associated with individual brain regions. Unlike previous models, each model only examined relationships within one whole brain area and did not calculate whole brain effects as an aggregate of individual regions.

Instead, the lowest level sampled β0_n_ and β1_n_ parameter posteriors at a singular volumetric area (whole brain, tissue, cerebrospinal fluid) for each subject n, and the second level constrained these distributions to be drawn from group-level Gaussian distributions, i.e., β0_n_∼N(μ_β0_,σ_β0_) and β1_n,r_∼N(μ_β1_,σ_β1_). As before, the group-level μ_β1_ parameter posterior reflects the expected value of the brain-hormone relation at the group level within the whole brain area, while σ_β1_ reflects variation in this relation across individuals at in that area, i.e., “shrinkage”. As before, we assessed relations by computing a deterministic posterior (D), which scaled the expected value by the shrinkage, i.e., μ_β1_/σ_β1_. Here, we imposed a region of practical equivalence of (-0.35 to 0.35) for D posteriors, determined with null-generative simulations using identical data structure (1 region, 30 participants, 3 sessions, variables drawn from a z-distribution).

## Acknowledgements

The authors thank Jan Martin and Frederik Bernd Lund at the Institute of Radiology, University Hospital Erlangen, Friedrich-Alexander-Universität Erlangen-Nürnberg (FAU), Erlangen, Germany for providing software code for the gradient tensor waveforms. We would also like to thank Mario Mendoza for MRI assistance, as well as Drs. Laura Pritschet, Shuying Yu, and Caitlin Taylor for advising on study procedures. The research was supported by the Institute for Collaborative Biotechnologies under Cooperative Agreement W911NF-19-2-0026 and contract W911NF-19-D-0001 from the Army Research Office. The content of the information does not necessarily reflect the position or the policy of the Government and no official endorsement should be inferred. For blood sample processing, the Endocrine Technologies Core (ETC) at Oregon National Primate Research Center (ONPRC) is supported (in part) by NIH grant P51 OD011092 for operation of the Oregon National Primate Research Center. Research reported in this publication was supported by the Office of the Director, National Institutes Of Health of the National Institutes of Health under Award Number S10OD026701. The content is solely the responsibility of the authors and does not necessarily represent the official views of the National Institutes of Health.

