## supplementary material for "Menstrual cycle-driven hormone concentrations co-fluctuate with white and grey matter architecture changes across the whole brain"

*Contributed equally as first author

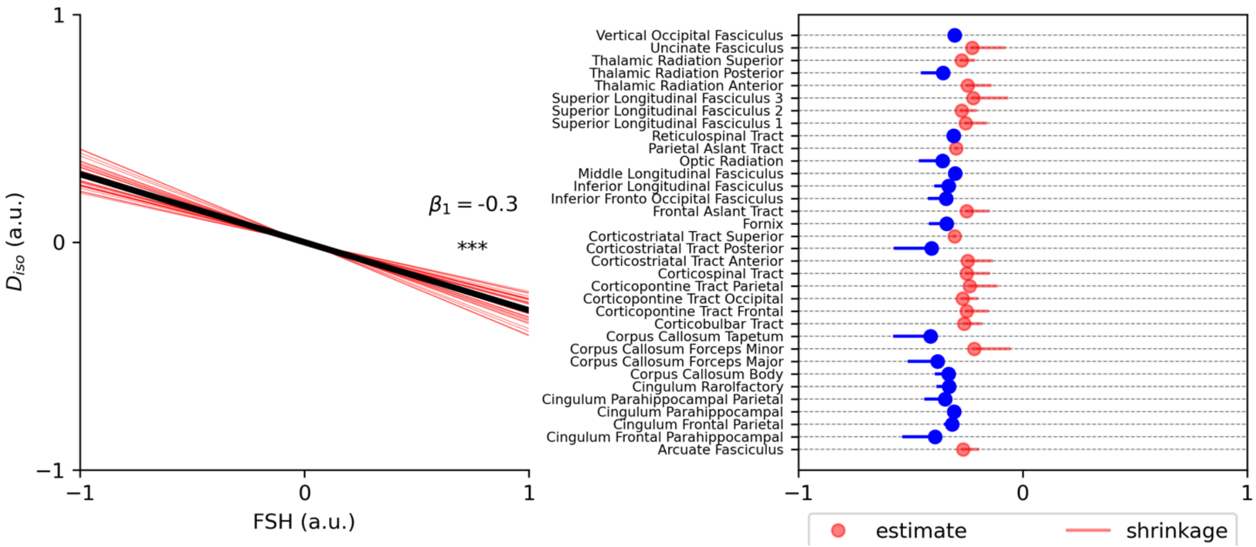


**FIGURE S1.** FSH-$D_{iso}$ whole brain and region-specific beta weight estimates as determined by hierarchical Bayesian regression modeling. Left panel: Representation of the FSH-$D_{iso}$ regression beta slope for whole brain (black line) and bilateral region (red lines) relationships. *** indicates that the HDI of D (whole brain effect posterior) exceeded a defined ROPE of -0.06 to 0.06, a.u. = arbitrary units. Right panel: Visualization of regression beta weight estimates and hierarchical Bayesian shrinkage (region-level estimates are affected by whole brain estimate) for each bilateral white matter HCP1065 Tractography Atlas ROI ($D_{iso}$ values were averaged between left and right hemispheres before entry into the model). Blue indicates that the HDI of $d_r$ for that region exceeded a defined ROPE of -0.35 to 0.35 (credible region effect). Red indicates that no credible effect was observed.

**TABLE S1.** Region-specific beta weights and credibility tests for FSH-$D_{iso}$ relationships

| Region | $\beta_1$ | D | HDI of D |
|---|---|---|---|
| Arcuate Fasciculus | -0.266 | -0.718 | [-1.252, -0.254] |
| **Cingulum Frontal Parahippocampal** | **-0.392*** | **-1.065** | **[-1.722, -0.545]** |
| **Cingulum Frontal Parietal** | **-0.316*** | **-0.831** | **[-1.379, -0.41]** |
| **Cingulum Parahippocampal** | **-0.306*** | **-0.801** | **[-1.315, -0.359]** |
| **Cingulum Parahippocampal Parietal** | **-0.348*** | **-0.907** | **[-1.453, -0.457]** |
| **Cingulum Rarolfactory** | **-0.328*** | **-0.878** | **[-1.44, -0.427]** |
| **Corpus Callosum Body** | **-0.331*** | **-0.902** | **[-1.49, -0.436]** |
| **Corpus Callosum Forceps Major** | **-0.380*** | **-1.113** | **[-1.934, -0.548]** |
| Corpus Callosum Forceps Minor | -0.216 | -0.450 | [-0.831, -0.084] |
| **Corpus Callosum Tapetum** | **-0.411*** | **-1.085** | **[-1.719, -0.569]** |
| Corticobulbar Tract | -0.262 | -0.650 | [-1.095, -0.223] |
| Corticopontine Tract Frontal | -0.251 | -0.630 | [-1.062, -0.198] |
| Corticopontine Tract Occipital | -0.268 | -0.690 | [-1.166, -0.249] |
| Corticopontine Tract Parietal | -0.237 | -0.588 | [-1.029, -0.162] |
| Corticospinal Tract | -0.251 | -0.649 | [-1.121, -0.203] |
| Corticostriatal Tract Anterior | -0.246 | -0.630 | [-1.106, -0.207] |
| **Corticostriatal Tract Posterior** | **-0.407*** | **-1.172** | **[-1.979, -0.585]** |
| Corticostriatal Tract Superior | -0.304 | -0.827 | [-1.374, -0.349] |
| **Fornix** | **-0.339*** | **-0.938** | **[-1.572, -0.454]** |
| Frontal Aslant Tract | -0.251 | -0.663 | [-1.148, -0.2] |
| **Inferior Fronto Occipital Fasciculus** | **-0.341*** | **-0.939** | **[-1.576, -0.475]** |
| **Inferior Longitudinal Fasciculus** | **-0.331*** | **-0.963** | **[-1.696, -0.453]** |
| **Middle Longitudinal Fasciculus** | **-0.301*** | **-0.812** | **[-1.358, -0.359]** |
| **Optic Radiation** | **-0.358*** | **-0.951** | **[-1.541, -0.484]** |
| Parietal Aslant Tract | -0.297 | -0.826 | [-1.409, -0.344] |
| **Reticulospinal Tract** | **-0.308*** | **-0.810** | **[-1.325, -0.368]** |
| Superior Longitudinal Fasciculus 1 | -0.254 | -0.664 | [-1.136, -0.219] |
| Superior Longitudinal Fasciculus 2 | -0.272 | -0.710 | [-1.199, -0.258] |
| Superior Longitudinal Fasciculus 3 | -0.222 | -0.567 | [-1.023, -0.102] |
| Thalamic Radiation Anterior | -0.247 | -0.593 | [-1.019, -0.196] |
| **Thalamic Radiation Posterior** | **-0.355*** | **-0.986** | **[-1.641, -0.491]** |
| Thalamic Radiation Superior | -0.272 | -0.728 | [-1.256, -0.28] |
| Uncinate Fasciculus | -0.225 | -0.524 | [-0.925, -0.117] |
| **Vertical Occipital Fasciculus** | **-0.304*** | **-0.857** | **[-1.479, -0.364]** |

* Indicates credible relation at the region level (HDI of $d_r$ lies outside of the ROPE [-0.35-0.35])

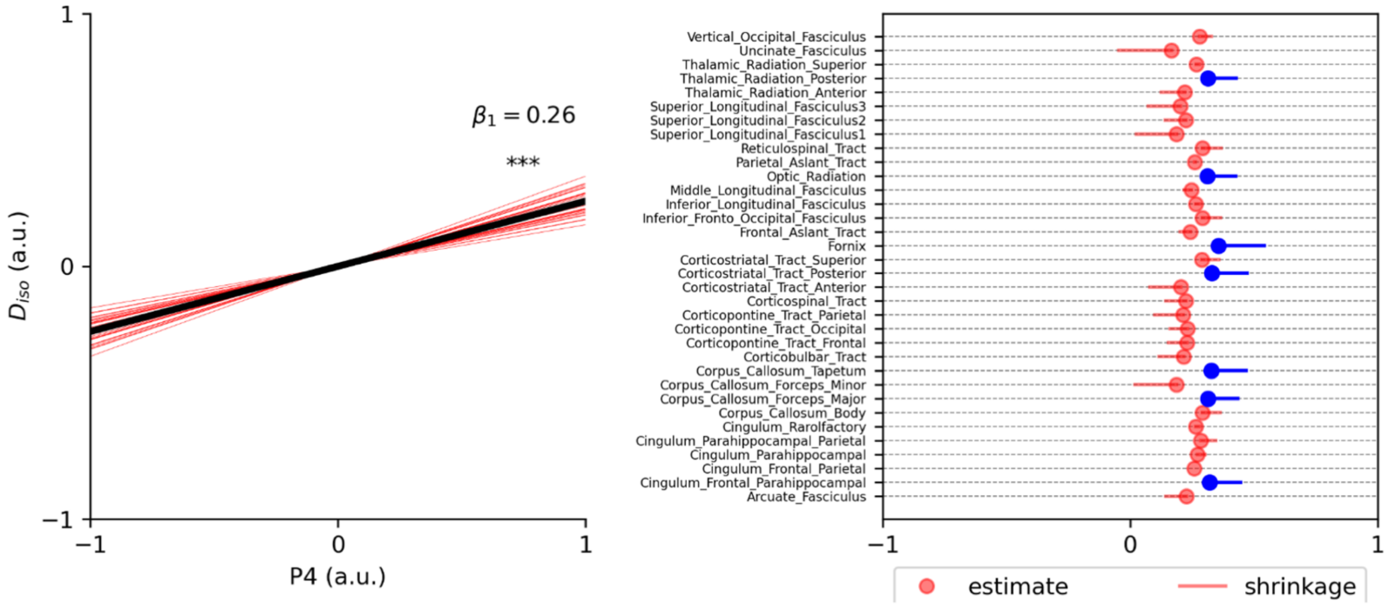


**FIGURE S2.** Progesterone-$D_{iso}$ whole brain and region-specific beta weight estimates as determined by hierarchical Bayesian regression modeling. Left panel: Representation of the progesterone-$D_{iso}$ regression beta slope for whole brain (black line) and bilateral region (red lines) relationships. *** indicates that the HDI of D (whole brain effect posterior) exceeded a defined ROPE of -0.06 to 0.06, a.u. = arbitrary units. Right panel: Visualization of regression beta weight estimates and hierarchical Bayesian shrinkage (region-level estimates are affected by whole brain estimate) for each bilateral white matter HCP1065 Tractography Atlas ROI ($D_{iso}$ values were averaged between left and right hemispheres before entry into the model). Blue indicates that the HDI of $d_r$ for that region exceeded a defined ROPE of -0.35 to 0.35 (credible region effect). Red indicates that no credible effect was observed.

**TABLE S2.** Region-specific beta weights and credibility tests for Progesterone-$D_{iso}$ relationships

| Region | $\beta_1$ | D | HDI of D |
|---|---|---|---|
| Arcuate Fasciculus | 0.225 | 0.523 | [0.137, 0.874] |
| Cingulum Frontal Parahippocampal | 0.321* | 0.738 | [0.393, 1.145] |
| Cingulum Frontal Parietal | 0.258 | 0.593 | [0.243, 0.949] |
| Cingulum Parahippocampal | 0.272 | 0.624 | [0.295, 1.002] |
| Cingulum Parahippocampal Parietal | 0.285 | 0.663 | [0.32, 1.054] |
| Cingulum Rarolfactory | 0.265 | 0.608 | [0.262, 0.971] |
| Corpus Callosum Body | 0.291 | 0.667 | [0.347, 1.066] |
| Corpus Callosum Forceps Major | 0.314* | 0.733 | [0.385, 1.155] |
| Corpus Callosum Forceps Minor | 0.185 | 0.410 | [0.036, 0.74] |
| Corpus Callosum Tapetum | 0.327* | 0.767 | [0.427, 1.227] |
| Corticobulbar Tract | 0.215 | 0.461 | [0.105, 0.762] |
| Corticopontine Tract Frontal | 0.227 | 0.502 | [0.15, 0.823] |
| Corticopontine Tract Occipital | 0.230 | 0.529 | [0.165, 0.875] |
| Corticopontine Tract Parietal | 0.212 | 0.478 | [0.105, 0.805] |
| Corticospinal Tract | 0.225 | 0.501 | [0.147, 0.825] |
| Corticostriatal Tract Anterior | 0.205 | 0.463 | [0.073, 0.787] |
| Corticostriatal Tract Posterior | 0.329* | 0.765 | [0.416, 1.205] |
| Corticostriatal Tract Superior | 0.289 | 0.672 | [0.329, 1.068] |
| Fornix | 0.357* | 0.836 | [0.464, 1.315] |
| Frontal Aslant Tract | 0.242 | 0.559 | [0.178, 0.91] |
| Inferior Fronto Occipital Fasciculus | 0.291 | 0.684 | [0.317, 1.077] |
| Inferior Longitudinal Fasciculus | 0.265 | 0.624 | [0.265, 1.012] |
| Middle Longitudinal Fasciculus | 0.247 | 0.562 | [0.208, 0.911] |
| Optic Radiation | 0.312* | 0.724 | [0.387, 1.139] |
| Parietal Aslant Tract | 0.260 | 0.596 | [0.246, 0.953] |
| Reticulospinal Tract | 0.291 | 0.685 | [0.349, 1.106] |
| Superior Longitudinal Fasciculus 1 | 0.186 | 0.410 | [0.032, 0.737] |
| Superior Longitudinal Fasciculus 2 | 0.224 | 0.511 | [0.147, 0.852] |
| Superior Longitudinal Fasciculus 3 | 0.203 | 0.456 | [0.073, 0.775] |
| Thalamic Radiation Anterior | 0.219 | 0.490 | [0.13, 0.813] |
| Thalamic Radiation Posterior | 0.313* | 0.721 | [0.39, 1.135] |
| Thalamic Radiation Superior | 0.266 | 0.618 | [0.273, 1.007] |
| Uncinate Fasciculus | 0.165 | 0.375 | [-0.047, 0.717] |
| Vertical Occipital Fasciculus | 0.279 | 0.645 | [0.312, 1.028] |

* Indicates credible relation at the region level (HDI of $d_r$ lies outside of the ROPE [-0.35-0.35])

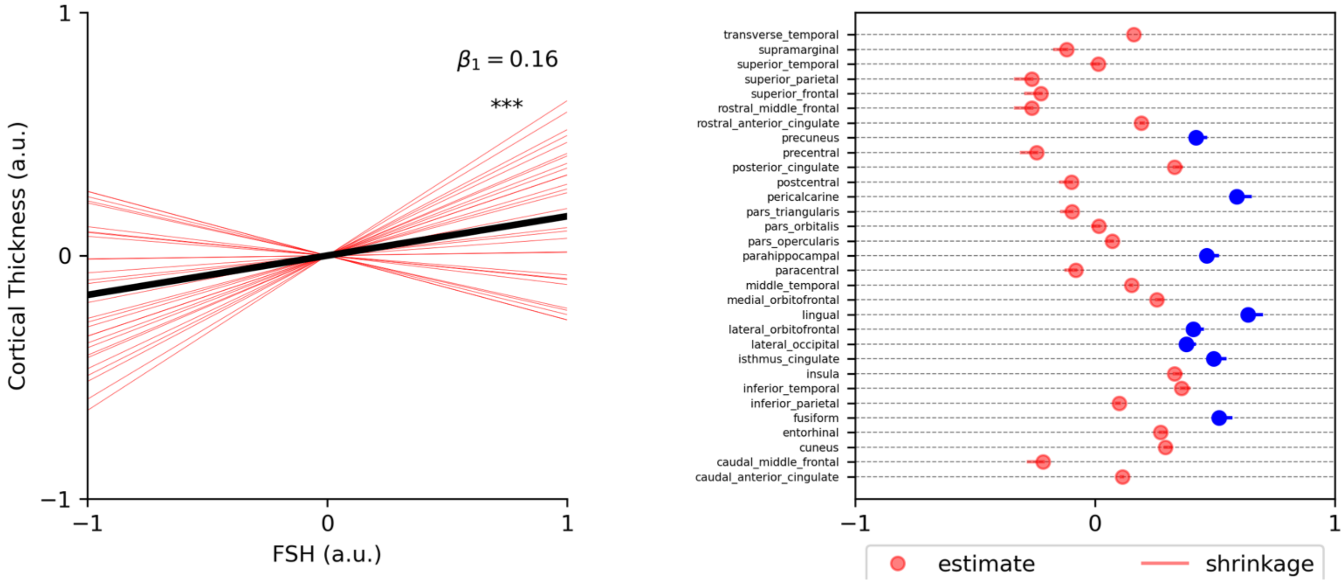


**FIGURE S3.** FSH-CT whole brain and region-specific beta weight estimates as determined by hierarchical Bayesian regression modeling. Left panel: Representation of the FSH-CT regression beta slope for whole brain (black line) and bilateral region (red lines) relationships. *** indicates that the HDI of D (whole brain effect posterior) exceeded a defined ROPE of -0.06 to 0.06, a.u. = arbitrary units. Right panel: Visualization of regression beta weight estimates and hierarchical Bayesian shrinkage (region-level estimates are affected by whole brain estimate) for each bilateral grey matter Desikan-Killiany-Tourville cortical ROI (CT values were averaged between left and right hemispheres before entry into the model). Blue indicates that the HDI of $d_r$ for that region exceeded a defined ROPE of -0.35 to 0.35 (credible region effect). Red indicates that no credible effect was observed.

**TABLE S3.** Region-specific beta weights and credibility tests for FSH-CT relationships

| Region | $\beta_1$ | D | HDI of D |
|---|---|---|---|
| Caudal Anterior Cingulate | 0.115 | 0.278 | [-0.253, 0.81] |
| Caudal Middle Frontal | -0.217 | -0.509 | [-1.039, 0.006] |
| Cuneus | 0.294 | 0.726 | [0.166, 1.3] |
| Entorhinal | 0.274 | 0.653 | [0.134, 1.196] |
| **Fusiform** | **0.518*** | **1.291** | **[0.699, 1.937]** |
| Inferior Parietal | 0.101 | 0.248 | [-0.304, 0.78] |
| Inferior Temporal | 0.360 | 0.887 | [0.333, 1.483] |
| Insula | 0.331 | 0.798 | [0.256, 1.358] |
| **Isthmus Cingulate** | **0.494*** | **1.248** | **[0.647, 1.921]** |
| **Lateral Occipital** | **0.379*** | **0.939** | **[0.376, 1.521]** |
| **Lateral Orbitofrontal** | **0.410*** | **1.009** | **[0.444, 1.618]** |
| **Lingual** | **0.637*** | **1.691** | **[0.979, 2.625]** |
| Medial Orbitofrontal | 0.259 | 0.655 | [0.09, 1.262] |
| Middle Temporal | 0.152 | 0.371 | [-0.153, 0.923] |
| Paracentral | -0.079 | -0.190 | [-0.717, 0.328] |
| **Parahippocampal** | **0.466*** | **1.139** | **[0.589, 1.754]** |
| Pars Opercularis | 0.071 | 0.173 | [-0.362, 0.715] |
| Pars Orbitalis | 0.015 | 0.035 | [-0.46, 0.534] |
| Pars Triangularis | -0.095 | -0.220 | [-0.741, 0.273] |
| **Pericalcarine** | **0.591*** | **1.568** | **[0.88, 2.467]** |
| Postcentral | -0.098 | -0.236 | [-0.783, 0.28] |
| Posterior Cingulate | 0.332 | 0.837 | [0.26, 1.458] |
| Precentral | -0.243 | -0.565 | [-1.08, -0.045] |
| **Precuneus** | **0.420*** | **1.041** | **[0.473, 1.648]** |
| Rostral Anterior Cingulate | 0.194 | 0.490 | [-0.061, 1.085] |
| Rostral Middle Frontal | -0.265 | -0.609 | [-1.152, -0.116] |
| Superior Frontal | -0.225 | -0.527 | [-1.074, -0.031] |
| Superior Parietal | -0.264 | -0.637 | [-1.196, -0.1] |
| Superior Temporal | 0.013 | 0.033 | [-0.531, 0.57] |
| Supramarginal | -0.119 | -0.28 | [-0.815, 0.229] |
| Transverse Temporal | 0.161 | 0.399 | [-0.142, 0.951] |

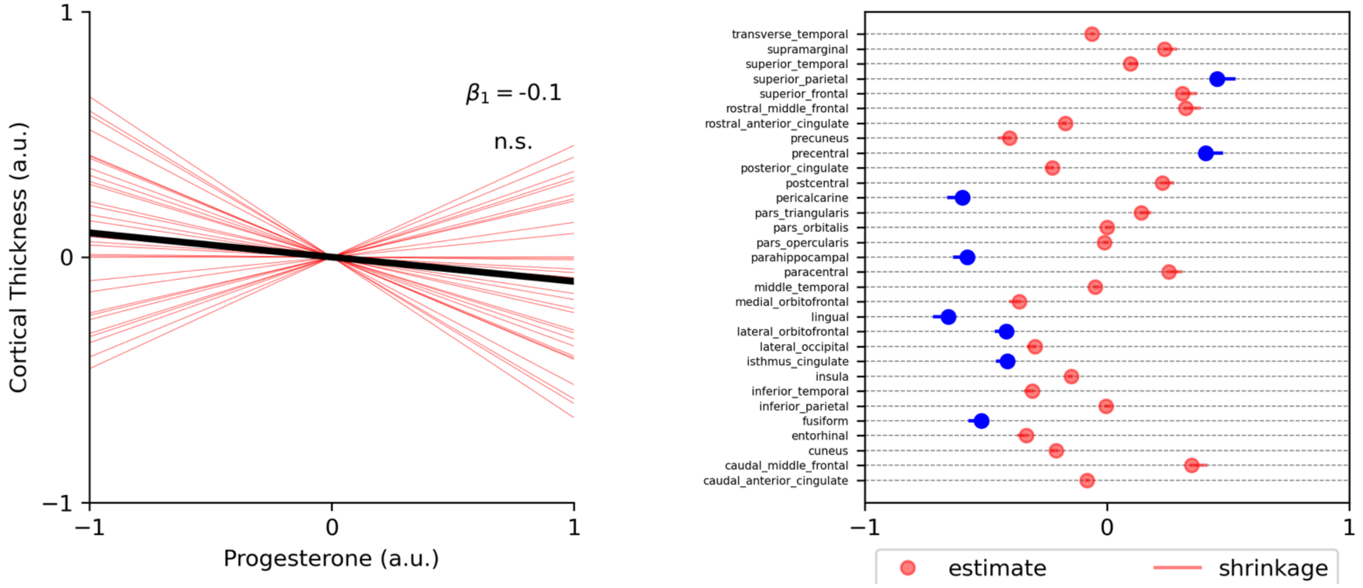


**FIGURE S4.** Progesterone-CT whole brain and region-specific beta weight estimates as determined by hierarchical Bayesian regression modeling. Left panel: Representation of the progesterone-CT regression beta slope for whole brain (black line) and bilateral region (red lines) relationships. *** indicates that the HDI of D (whole brain effect posterior) exceeded a defined ROPE of -0.06 to 0.06, a.u. = arbitrary units. Right panel: Visualization of regression beta weight estimates and hierarchical Bayesian shrinkage (region-level estimates are affected by whole brain estimate) for each individual grey matter Desikan-Killiany-Tourville cortical ROI (CT values were averaged between left and right hemispheres before entry into the model). Blue indicates that the HDI of $d_r$ for that region exceeded a defined ROPE of -0.35 to 0.35 (credible region effect). Red indicates that no credible effect was observed.

**TABLE S4.** Region-specific beta weights and credibility tests for Progesterone-CT relationships

| Region | $\beta_1$ | D | HDI of D |
|---|---|---|---|
| Caudal Anterior Cingulate | -0.083 | -0.167 | [-0.647, 0.304] |
| Caudal Middle Frontal | 0.349 | 0.699 | [0.248, 1.18] |
| Cuneus | -0.210 | -0.414 | [-0.868, 0.037] |
| Entorhinal | -0.334 | -0.661 | [-1.112, -0.203] |
| **Fusiform** | **-0.52*** | **-1.027** | **[-1.485, -0.589]** |
| Inferior Parietal | -0.005 | -0.009 | [-0.467, 0.427] |
| Inferior Temporal | -0.309 | -0.608 | [-1.061, -0.161] |
| Insula | -0.148 | -0.289 | [-0.728, 0.162] |
| **Isthmus Cingulate** | **-0.412*** | **-0.823** | **[-1.287, -0.369]** |
| Lateral Occipital | -0.297 | -0.594 | [-1.061, -0.129] |
| **Lateral Orbitofrontal** | **-0.416*** | **-0.810** | **[-1.256, -0.384]** |
| **Lingual** | **-0.654*** | **-1.365** | **[-1.921, -0.866]** |
| Medial Orbitofrontal | -0.362 | -0.743 | [-1.235, -0.253] |
| Middle Temporal | -0.049 | -0.097 | [-0.551, 0.378] |
| Paracentral | 0.255 | 0.508 | [0.056, 0.971] |
| **Parahippocampal** | **-0.578*** | **-1.141** | **[-1.596, -0.716]** |
| Pars Opercularis | -0.011 | -0.021 | [-0.481, 0.415] |
| Pars Orbitalis | 0.001 | 0.001 | [-0.451, 0.44] |
| Pars Triangularis | 0.142 | 0.283 | [-0.175, 0.756] |
| **Pericalcarine** | **-0.596*** | **-1.24** | **[-1.799, -0.746]** |
| Postcentral | 0.228 | 0.454 | [0.003, 0.926] |
| Posterior Cingulate | -0.226 | -0.445 | [-0.894, 0.009] |
| **Precentral** | **0.407*** | **0.823** | **[0.361, 1.311]** |
| Precuneus | -0.403 | -0.820 | [-1.3, -0.337] |
| Rostral Anterior Cingulate | -0.173 | -0.347 | [-0.829, 0.111] |
| Rostral Middle Frontal | 0.325 | 0.651 | [0.185, 1.121] |
| Superior Frontal | 0.311 | 0.628 | [0.161, 1.099] |
| **Superior Parietal** | **0.455*** | **0.926** | **[0.459, 1.432]** |
| Superior Temporal | 0.096 | 0.189 | [-0.254, 0.633] |
| Supramarginal | 0.237 | 0.469 | [0.027, 0.938] |
| Transverse Temporal | -0.063 | -0.125 | [-0.583, 0.33] |

*Supplementary Methods: Determination of Region of Practical Equivalence (ROPE) for Credibility Testing*

To focus our discussions on only robust effects from our hierarchical Bayes models, we implemented a policy whereby we only considered a relation credible if the highest density interval (HDI) of the shrinkage-adjusted expected value of its coefficient (i.e., $d_r$ and D, as described in the methods) not only fell above or below zero, but additionally fell above or below a region of practical equivalence (ROPE) that we estimated from null-generative simulations.

Note that in our models, we z-score both the independent and dependent variables (each 3-element vectors) at the lowest level of the hierarchy. All low-level regression coefficients are therefore mapping the relation between two three-element vectors drawn from a z-distribution. We accordingly ran repeated simulations, where we computed coefficients mapping the relation between two 3-element vectors drawn randomly from z-distributions to see what range of values emerge with different data structures in a null-generative context.

More specifically, for relations in the regionalized models (i.e., using the $d_r$ metric described in the methods), we computed a ROPE with the following: on each of 100,000 simulations (k), we computed 30 (i.e., n = 30 participants) regression coefficients ($\beta 1_{k,n}$), mapping $x_{k,n}$ and $y_{k,n}$ with general linear model formula $y_{k,n} = \beta 1_{k,n} x_{k,n} + c$. $x_{k,n}$ and $y_{k,n}$ were both three-element vectors drawn from the z-distribution, i.e., $[x_{k,n}, y_{k,n}] \sim N(0,1)$. We computed an equivalent $\mu_{\beta 1(k)}$ for each simulation by averaging the 30 (i.e., n) $\beta 1_k$ estimates, i.e.,

$$\mu_{\beta 1(k)} = 1/n \sum_{i=1}^{n} \beta 1_{k,n}$$

and an equivalent $\sigma_{\beta 1(k)}$ by computing their standard deviation, i.e.,

$$\sigma_{\beta 1(k)} = \sqrt{\frac{\sum_{i=1}^{n} \left(\beta 1_{k,n} - \mu_{\beta 1(k)}\right)^2}{n-1}}$$

We finally computed a k-element distribution of d values, i.e., where $d_k = \mu_{\beta 1(k)} / \sigma_{\beta 1(k)}$. 94% of the null-generative simulations of d fell between -0.35 and 0.35, which we used as the boundaries of our ROPE.

For whole-brain relations in regionalized brain models (i.e., using the D metric described in the methods), we computed a ROPE with the following: on each of 100,000 simulations (k), we computed 34 (diffusion measures) or 31 (cortical thickness) regions (r) of 30 (i.e., n = 30 participants) regression coefficients ($\beta 1_{k,n,r}$), mapping $x_{k,n,r}$ and $y_{k,n,r}$ with general linear model formula $y_{k,n,r} = \beta 1_{k,n,r} x_{k,n,r} + c$. $x_{k,n,r}$ and $y_{k,n,r}$ were both three-element vectors drawn from the z-distribution, i.e., $[x_{k,n,r}, y_{k,n,r}] \sim N(0,1)$. We computed an equivalent $\mu_{\beta 1(k,r)}$ for each region in each simulation by averaging the 30 (i.e., n) $\beta 1_{k,r}$ estimates, i.e.,

$$\mu_{\beta 1(k,r)} = 1/n \sum_{i=1}^{n} \beta 1_{k,n,r}$$

and a variance estimate $\sigma^2_{\beta 1(k,r)}$ by computing their variance, i.e.,

$$\sigma^2_{\beta 1(k,r)} = \frac{\sum_{i=1}^{n} (\beta 1_{k,n,r} - \mu_{\beta 1(k,r)})^2}{n-1}$$

Then, across regions on each simulation, we computed the equivalent of $M_{u(\beta 1)}$ with the average across $\mu_{\beta 1(k,r)}$ values, i.e.,

$$M_{u(\beta 1,k)} = 1/r \sum_{i=1}^{r} \mu_{\beta 1(k,r)}$$

and $M_{\sigma(\beta 1)}$ with the root mean of $\sigma^2_{\beta 1(k,r)}$ values, i.e.,

$$M_{\sigma(\beta 1,k)} = \sqrt{1/r \sum_{i=1}^{r} \sigma^2_{\beta 1(k,r)}}$$

We finally computed a k-element distribution of D values, i.e., where $D_k = M_{u(\beta 1,k)} / M_{\sigma(\beta 1,k)}$. 94% of the null-generative simulations of D fell between -0.06 and 0.06, which we used as the boundaries of our ROPE.
